# Can the occipital alpha-phase speed up visual detection through a real-time EEG-based brain-computer interface (BCI)?

**DOI:** 10.1101/2020.07.06.189712

**Authors:** Irene Vigué-Guix, Luis Morís Fernández, Mireia Torralba Cuello, Manuela Ruzzoli, Salvador Soto-Faraco

**Affiliations:** Center for Brain and Cognition, Departament de Tecnologies de la Informació i les Comunicacions, Universitat Pompeu Fabra, 08005, Barcelona, Spain; Departamento de Psicología Básica, Universidad Autónoma de Madrid, Madrid, Spain; Institute of Neuroscience and Psychology, University of Glasgow, G12 8QB, Glasgow, United Kingdom; Institució Catalana de Recerca i Estudis Avançats (ICREA), 08010, Barcelona, Spain

**Keywords:** EEG, oscillations, alpha, phase, real-time, brain-computer interface (BCI), response time, visual perception

## Abstract

Electrical brain oscillations reflect fluctuations in neural excitability. Fluctuations in the alpha band (α, 8-12 Hz) in the occipito-parietal cortex are thought to regulate sensory responses, leading to cyclic variations in visual perception. Inspired by this theory, some past and recent studies have addressed the relationship between α-phase from extra-cranial EEG and behavioural responses to visual stimuli in humans. The latest studies have used *offline* approaches to confirm α-gated cyclic patterns. However, a particularly relevant implication is the possibility to use this principle *online* for real-time neurotechnology, whereby stimuli are time-locked to specific α-phases leading to predictable outcomes in performance. Here we aimed at providing a proof-of-concept for such real-time neurotechnology. Participants performed a speeded response task to visual targets that were presented upon a real-time estimation of the α-phase via an EEG closed-loop brain-computer interface (BCI). We predicted, according to the theory, a modulation of reaction times (RTs) along the α-cycle. Our BCI system achieved reliable trial-to-trial phase-locking of stimuli to the phase of individual occipito-parietal α-oscillations. Yet, the behavioural results did not support a consistent relation between RTs and the phase of the α-cycle neither at group nor single participant levels. We must conclude that although the α-phase might play a role in perceptual decisions from a theoretical perspective, its impact on EEG-based BCI application appears negligible.

## INTRODUCTION

Alpha oscillations (α, 8-12 Hz) in the occipito-parietal cortex reflect ongoing fluctuations in cortical excitability (Bishop, 1932; Adrian & Matthews, 1934; Worden *et al.*, 2000; Kelly *et al.*, 2006). It follows that the perceptual fate of a visual stimulus would depend upon the instant it evokes neural activity within the ongoing α-cycle, leading to cyclic alternations between more and less favourable phases for perceptual processing (Klimesch *et al.*, 2007; Jensen & Mazaheri, 2010; Klimesch, 2012; Jensen *et al.*, 2014; VanRullen, 2016a). This hypothesis has been entertained for nearly a hundred years (since Bishop, 1932), and it has revived recently (see VanRullen, 2016a). The fact that α-fluctuations can be picked up extra-cranially via magnetic- or electrical encephalography (M/EEG) makes α an optimal candidate to study human perception non-invasively. Specifically, both the power (Worden *et al.*, 2000; Ergenoglu *et al.*, 2004; Babiloni *et al.*, 2006; Kelly *et al.*, 2006; Thut *et al.*, 2006; Klimesch *et al.*, 2007; Palva & Palva, 2007; Foxe & Snyder, 2011) and the phase of the occipito-parietal α (Klimesch *et al.*, 2007; Palva & Palva, 2007; Mathewson *et al.*, 2009; Klimesch, 2012; Jensen *et al.*, 2014; VanRullen, 2016b) have been linked to performance in visual perception (van Dijk *et al.*, 2008; Jensen & Mazaheri, 2010; Jensen *et al.*, 2011; Samaha & Postle, 2015; VanRullen, 2016a). However, while the evidence for the role of α-power in perceptual judgments seems well-established (Walsh, 1952; Lansing *et al.*, 1959; Jensen *et al.*, 2011; Bompas *et al.*, 2015; but see Benwell *et al.*, 2017), the role of the α-phase is still not clearly settled (Walsh, 1952; O’Hare, 1954; Benwell *et al.*, 2017; Ruzzoli *et al.*, 2019).

So far, most studies have used an *offline* approach to study changes in perception based on the α-phase. In these studies, stimuli that demand a behavioural response are presented at random (or pseudo-random) intervals while EEG is recorded, and at a later time trials are separated in terms of the behavioural outcome (hit vs miss; fast vs slow reaction times-RTs) and sorted post-hoc based on which phase within the pre-target α-cycle, the stimuli happened to fall. Average responses are then statistically compared across phase bins to conclude on a phase-behavioural relationship (see Busch *et al.*, 2009 for an example). A significant correlation between phase and behaviour, when found, provides support for the theory.

However, the possibility to link α-fluctuations to behaviour via non-invasive methods, such as EEG, is also attractive given the potential applications in Brain-Computer Interfaces (BCI) (Jensen *et al.*, 2011; Zrenner *et al.*, 2016). For example, one could design closed-loop BCI systems that deliver information at favourable brain states for perceptual encoding to improve alerting, learning or memory (Brunner *et al.*, 2015; see Zrenner *et al.*, 2016 for examples in the motor domain).

To harness on the α-theories to develop BCI systems, one must use an *online* approach, which should be efficient even at the single-subject level. Real-time EEG analysis allows BCI settings to trigger stimuli at precise phase angles during the ongoing fluctuations in the individual α-rhythm that are thought to be associated with specific outcomes (hit/miss, fast/slow RTs). This approach exploits a specific brain-behaviour relationship (e.g., α-phase and perception) to augment encoding of information with millisecond precision. The efficiency of closed-loop BCI must rely on predictable brain-behaviour relationships, in which the relevant parameters at play must be known beforehand. Hence, in turn, the attempt at using a closed-loop BCI approach is a test bench of neuro-cognitive theories such as the α-theories.

Interestingly, a good number of studies in the sixties already capitalized on the idea of time-locking stimulus presentation to the α-phase in real-time. Such attempts were popular enough by the middle of the decade as to prompt Callaway & Layne to write “*The idea of presenting photic stimuli at various phases of the spontaneous alpha rhythm to alter degrees of photic driving has occurred to many investigators*” (Callaway & Layne, 1964, pp 421). In one preeminent study published in the journal *Science* in 1960, Callaway & Yeager (Callaway & Yeager, 1960) endeavoured a closed-loop BCI addressing the relationship between the α-phase and RTs to visual events. They found that RTs were modulated as a function of the instant within the α-cycle the target flash was presented (see Lansing, 1957; Dustman & Beck, 1965 for similar results also in a real-time setting). Despite the substantial potential impact such closed-loop BCI on both theory and application, to the best of our knowledge, no modern study has implemented and reported a similar real-time protocol harnessing on occipital α-phase. The 60-year hiatus is remarkable, especially considering the α-theories are still entirely current up to this date.

Here, we capitalized on the putative relationship between the phase of ongoing α-oscillations and visual perception adopting a closed-loop BCI approach to provide new evidence for the α-theories, which opened enduring questions a long time ago. At the same time, the present study aimed at providing a proof of concept for the use of the phase of ongoing α-oscillations as a control signal in a closed-loop BCI system for practical applications. This experiment is a modern replication of Callaway & Yeager’ study (1960). We employed a visual speeded detection task in which the visual target was triggered in real-time as a function of the phase of the participant’s α-cycle. We expected that visual RTs would fluctuate along the α-cycle, or at least, that it should be possible to find two distinct phases associated with fast and slow RTs, respectively. The hypothesis, the procedure and the pipeline of the analysis were pre-registered before data collection (https://osf.io/nfdsv/). Deviations from the pre-registered procedure and exploratory analyses are clearly stated in the manuscript.

## METHODS

### Participants

#### Sample size

We planned a maximum sample of 16 participants with a stopping rule set after a minimum of 8 participants (see details below). Participants were selected without previous history of neurological or psychiatric diseases, with normal or corrected to normal vision, within 18-35 years old. The minimum/maximum sample size was decided a priori based on a Monte Carlo simulation on Callaway & Yeager’ data (1960) (See Figure S1). We estimated that if less than 3 participants out of 8 showed a significant difference between fast and slow phase bins, then the size of the effect in this experiment would be null or negligible compared to the original study (Callaway & Yeager, 1960), assuming an error of 5%.

#### Exclusion criteria

A participant was excluded if any of the following criteria were met: (1) No peak within the α-band: This criterion applied to the screening stage and ensured that the individual’s endogenous α*-*oscillation could be registered with a sufficiently high signal-to-noise ratio (SNR) to enable the BCI system to estimate instantaneous phases from the EEG signal reliably. This decision was based on two sub-criteria: strength and uniqueness (see *Screening and estimation of the Individual Frequency of Interest* section for more details). (2) Experiment duration: Given that we had a block stopping rule based on the number of trials per phase bin, the length of the experiment could vary as a function of how frequently the EEG phase could be reliably estimated for stimulus presentation. Hence, we had to establish an experiment duration limit. We decided to stop the experiment if a participant spent more than 10 minutes in the training block or two consecutive blocks within the real-time experimental stages. This criterion was added after we had run the first 2 participants, which required an update of the pre-registered protocol^1^.

We recruited 27 participants, 6 of which were discarded for not satisfying the required α-peak criterion in the screening stage, and 13 because of the duration criterion. The remaining 8 participants (aged 19-30 years, average 24 years, three females; all right-handed) completed the experiment. Data from excluded participants were not analysed. All the participants took part in the study voluntarily after giving informed consent, and they were compensated for their time with 10€ per hour. The duration of the experiment varied between 70 and 120 min. The study was designed in accordance with the Declaration of Helsinki and approved by the local ethics committee CEIC Parc de Mar (University Pompeu Fabra, Barcelona, Spain) before starting the recruitment.

### Experimental procedure

The experimental protocol started with a screening, followed by a training and two consecutive experimental stages (explained below). In the training and experimental stages, participants performed a speeded visual detection task in which stimuli were presented according to the phase of the individual spontaneous α-activity in real-time (see Figure 1).

**Figure 1.**
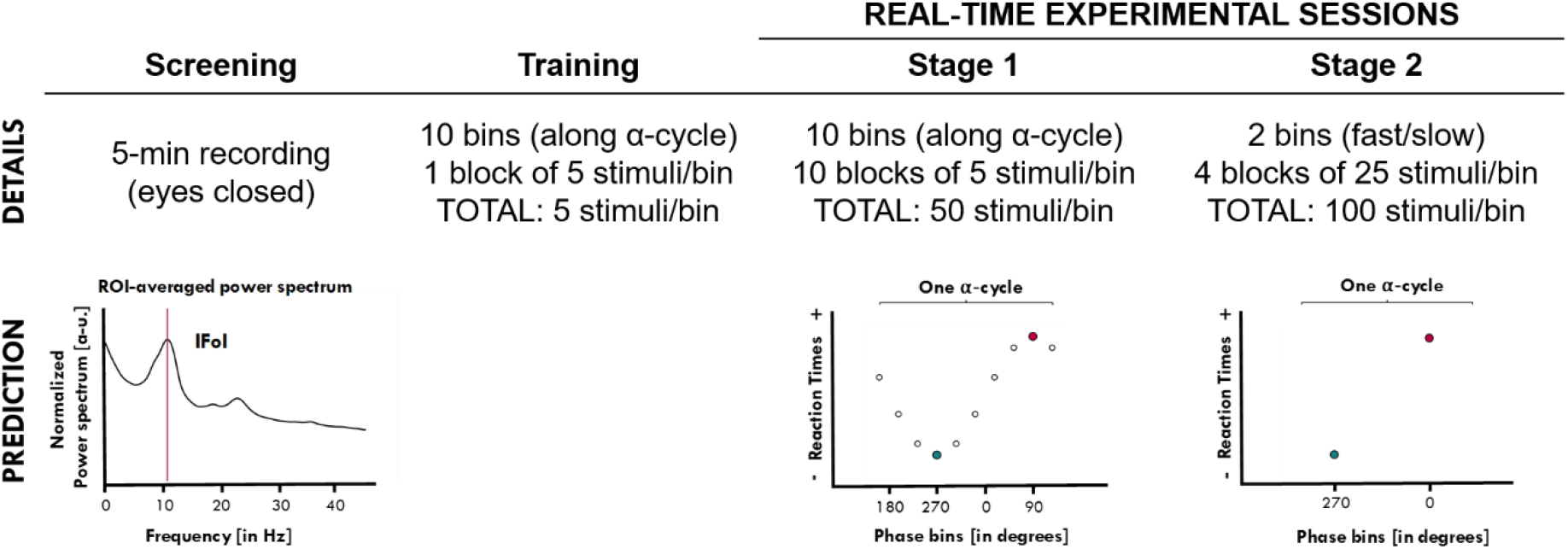
The protocol included four stages: Screening, Training, Stage 1 and Stage 2 of testing. During the Screening, we looked for the individual frequency of interest (IFoI) over the occipito-parietal area in a 5 min EEG recording at rest (eyes closed). Participants who did not display a single peak in the EEG within the range of interest (5-15 Hz) did not proceed to the following parts of the protocol. Included participants performed a Training, and entered Stage 1 of the real-time experimental sessions. In Stage 1, visual stimuli were triggered phase-locked to 10 equally spaced phase bins along the IFoI-cycle. For illustration purposes of an ideal theoretical outcome, stimulus onset (circles) is represented as a function of the EEG alpha cycle through a cosine wave. From Stage 1, the specific phase bins associated with the faster (green dot) and slower (red dot) RTs were selected for each participant and used for Stage 2. In Stage 2, visual stimuli were triggered only at the two-phase bins (fast/slow) individually selected from Stage 1. As illustrated, we predicted that fast and slow phase bins would lead to fast and slow RTs, respectively.

#### Task

Participants sat on a comfortable chair wearing an EEG cap, and a pair of opaque sunglasses, with two LEDs, mounted on each lens. The LEDs were controlled through the parallel port (both LEDs switched on and off simultaneously; luminance 0.076 cd/m^2^ at an approximate distance to participants’ eye of 1 cm)^2^. Participants were asked to keep their eyes closed throughout the experiment to maintain α-activity high and to limit eye movements (a strategy first suggested by Callaway & Yeager, 1960). We instructed the participants to remain attentive and to respond to the visual flashes as fast as possible by pressing a button in a response box using their right index finger. After the response (or after 1 second time out), the LEDs were switched off. An inter-trial interval (ITI), randomly chosen between 1500 and 2500 ms, was introduced between the button response (or 1s time out) and the beginning of the next trial. RTs were measured from the onset of the visual target until a button press was detected.

#### Training stage

Before the real-time experimental stages, participants were familiarised with the task in a training block (50 trials), identical to Stage 1 (see below).

#### Experimental stages

After training, participants went on to Stage 1, where visual targets were aimed at 10 equally-spaced phase bins covering the whole α-cycle (see *Real-time stimulus presentation*). This experimental stage was divided into 10 blocks, each ending after the acquisition of at least 5 valid RTs per phase bin, for a total of 50 RTs per phase bin across blocks. Once Stage 1 had been completed, the phase-bins corresponding to the fastest and slowest mean RTs were selected and used for Stage 2 (at the individual level). No statistical test was performed at Stage 1. Stage 2 started right after Stage 1. In Stage 2, visual targets were aimed only at the “fast” and the “slow” phase bins, estimated from Stage 1. Participants ran 4 blocks, each block ended after the collection of at least 25 valid trials for each of the two-phase bins, for a minimum total of 100 trials per bin.

#### EEG recording

Continuous EEG data were recorded at 500 Hz using the ENOBIO 20 5G system (Neuroelectrics, Barcelona, Spain) from 14-channels (F3, Fz, F4, C3, Cz, C4, P7, P3, Pz, P4, P8, O1, Oz, O2) with Cl-Ag electrodes placed according to the 10-20 international system. Impedance was kept below 10 kOhm, according to the Enobio coded system. Additional external electrodes were used to record vertical and horizontal eye movements. Electrode AFz was used as online reference and the right mastoid as ground. Activity from the left mastoid was recorded for offline re-referencing.

#### Screening and estimation of the Individual Frequency of Interest (IFoI)

We recorded 5 minutes of resting EEG with the eyes closed, which we used to determine the specific frequency of interest within the α-band for the real-time stages of the experiment. We estimated the power spectrum density (PSD) within the α-band (5-15 Hz) over occipito-parietal electrodes (OP-cluster: P7, P3, Pz, P4, P8, O1, Oz, O2) using the Welch method (window = 500 ms; overlap = 10%; resolution = 0.25 Hz). For each participant, the power spectrum was averaged across the electrodes of interest and normalized by the mean power spectrum from 1 to 40 Hz. We verified the strength of the peak - power at the local maximum within the 5-15 Hz window is greater than average power in the 1-40 Hz window - and its uniqueness - the peak is a single local maximum within a ± 5 Hz band. If a single frequency peak existed, it was considered as the IFoI^3^ and used later as a parameter for real-time analyses (Figure 1). If a unique frequency peak could not be detected, the participant was excluded from the study (see *Exclusion Criterion 1*).

#### Real-time stimulus presentation

We developed a BCI setting to trigger flashes (LEDs) at a specific α-phase based on real-time data from electrode O1 (as in Callaway & Yeager, 1960) through custom-built code in MATLAB (MathWorks, R2015.b). We used the *Lab Streaming Layer (LSL)* library (Swartz Center for Computational Neuroscience, UCSD, January 2018) to acquire EEG data with the ENOBIO acquisition software (NIC V2.0). Given that synchronization between the EEG time acquisition and local PC time is not supported for online streams by ENOBIO, we used an external signal (a parallel port pin connected to one of the ENOBIO electrodes) as time reference. In each iteration, we randomly pre-selected, among the bins available (10 bins in Stage 1, or 2 bins in Stage 2), a phase bin when to send the stimulus trigger. Ten-seconds of EEG data from the O1-electrode were continuously buffered, demeaned, and band-pass filtered (Butterworth forward filter, order 2, IFoI ± 5 Hz). Amplitude and phase were estimated at each time point using the Hilbert-transform.

When the average amplitude in the last second of the buffer was above 30% of the median value of the buffered data, a reference time point was set at the peak (90°) of the last α-cycle. Then, the moment at which the phase of interest in a given trial would occur was forecasted by extending a sine wave (frequency = IFoI) from the reference point. We targeted one of the 10 phase bins (36° each) within the next α-cycle starting after a safety buffer of 72° (~20 ms) for computation time. The stimulus trigger switched on the LEDs at the latency corresponding to the centre of the targeted phase bin (error ≤ 2 ms).

#### Responses and trial selection

Responses were collected from a button press via a response box connected to the parallel port. There was a response time out of 1 second after the stimulus, following which the next trial iteration began. If a response was registered before time out, we stored the RT and checked the accuracy of the real-time phase estimation by calculating the difference between the empirical phase at which the stimulus was delivered (according to the recorded EEG) and the intended one, using the Circular Statistics Toolbox in MATLAB (Berens, 2009). In Stage 1, if the stimulus had been triggered in an unintended phase, the trial was relocated to the actual (empirically measured) phase bin. In Stage 2, we only targeted two phases and set a tolerance of ± 1 phase bin to reduce the testing time, and we did not relocate trials. A trial was excluded if the empirical phase did not fall within the tolerance zone, or it fell in an overlapping bin between the slow and the fast bin (this could happen if the fast/slow bins were less than 72° apart).

Only trials that satisfied all the following criteria were accepted as valid: (1) Reaction time criterion: RTs within 50 and 300 ms (as in Callaway & Yeager, 1960). (2) Amplitude criterion: the amplitude of the IFoI-cycle window centred at stimulus onset had to be above the 30% of the median amplitude in the last 10s (this threshold criterion facilitated reliable phase estimation at stimulus presentation).

A block stopped when the intended number of trials per phase bin was reached (N=5 in Stage 1 or N=25 in Stage 2). In between blocks, participants took a break before starting a new one. Excess trials (which could happen due to trial relocation) were discarded.

### Statistical analyses

Stage 1 was designed to estimate the phase bins corresponding to faster and slower RTs throughout the α-phase, whereas Stage 2 provided data for the validation of the hypothesis. In Stage 1, we only ran descriptive statistics to calculate average RTs per phase bin, and to select the phase bins of interest. If the fast and slow phase bins selected in Stage 1 would indeed be representative of neural excitability states related to visual perception, then sending targets to these phase bins in Stage 2 should induce faster and slower responses, respectively. To test this prediction, in Stage 2, we performed individual and group-level analyses. For the individual analysis, we assessed the difference between the RTs collected in the predicted slow and predicted fast phase bins by a one-tailed t-test (independent samples) with α-level = 0.05. Note that in Stage 2, if slow/fast bins (± 1 phase bin) shared a common bin, then trials in that bin were post-hoc excluded and not used for the analysis. For the group analysis, we evaluated the difference between the mean RTs collected in the slow vs fast phase bins across participants by a one-tailed paired t-test with α-level = 0.05.

## RESULTS

Here, we present the results obtained from the pre-registered analyses as explained above, which replicated the conditions of the seminal study by Callaway & Yeager (1960), followed by reality checks and a set of exploratory analyses.

### Results of the pre-registered analyses

#### Stage 1: Selection of the fast/slow phase bins along α-cycle

We collected an average of 1034 (SD=219) responses per participant. An average of 77 (SD=53; 7.41%) trials were excluded because the RTs fell out of the 50 – 300 ms range and 341 (SD=190; 33%) trials were excluded because the amplitude threshold criterion was not met, leaving an average of 617 (SD=36; 60%) valid trials per participant. Among the valid trials, 280 (SD=35; 45.33%) trials hit the target phase bin of interest whereas 337 (SD=61; 54.67%) trials had to be relocated to the intended phase bin offline (most of them fell on neighbouring bins, see *Reality check 1: Accuracy of phase estimation during the real-time experiment*, below). Therefore, we reached the intended 50 valid trials per bin for each participant (following the elimination of excess trials). RTs for valid trials were on average 206 ms (SD=9 ms). We calculated the mean RT for each phase bin along the α-cycle for each participant (see Figure 2 and Table S1; see Table S2 for information about the number of trials at the individual level), and selected the phase bins associated with the slowest and fastest mean RTs, to be used in Stage 2. Overall, the mean RT difference between slow and fast phase bins in Stage 1 was 12 ms (SD=4; Max=17 ms; Min=8 ms).

**Figure 2.**
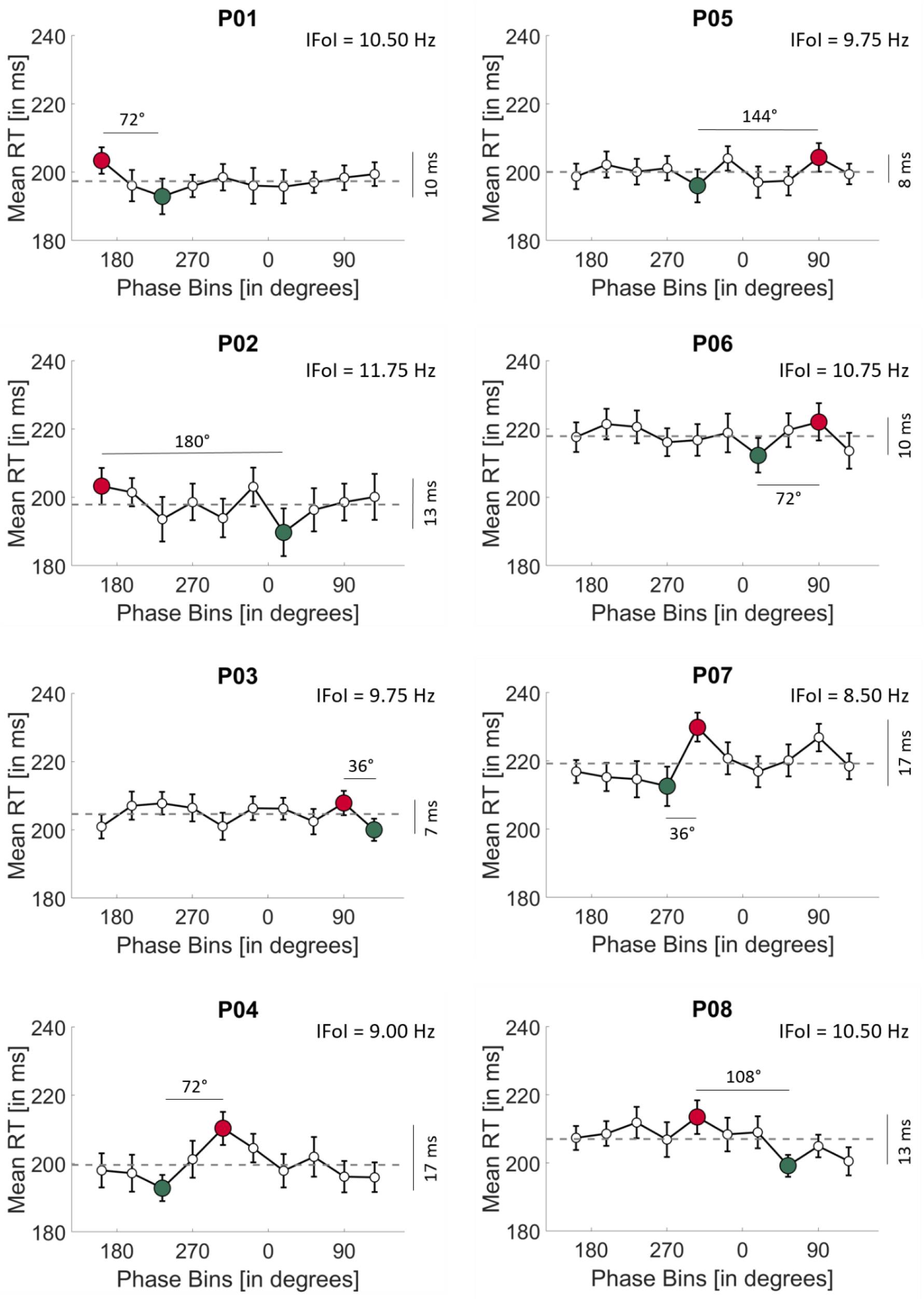
Individual mean RT (in ms, y-axis) plotted against the 10 phase bins (in degrees, x-axis) tested in Stage 1. The horizontal dashed line indicates the individual mean RT across all bins. For each participant, the graphs report the angular difference and the RT difference between the fast (green dot) and slow (red dot) phase bins. IFoI= Individual Frequency of Interest; Error bars = Standard Error of the Mean.

At this point, if the distribution of slow and fast RTs meets the expectations of the α-theory, the corresponding slow and fast phase bins should fall on roughly opposite angles (approximately 180°). However, what we observed is that for most of the participants, slow and fast phase bins were closer than 180°, being the mean difference 90° (SD=51°).

#### Stage 2: Validation of the α-phase relation to RT speed

In Stage 2, we collected an average of 408 (SD=103) trials per participant. Among these, 31 (SD=14; 7.65%) trials were excluded because they fell outside the RT criterion, 144 (SD=84; 35.30%) trials were excluded for not satisfying the amplitude threshold criterion, and 33 (SD=24; 8.09%) trials for not falling in the bin acceptance zone. After trial exclusion, we were left with a total of 200 valid trials each participant (100 trials per bin), as intended. Among these, an average of 107 (SD=24; 54%) trials hit the target phase bin of interest (± 1 bin), whereas 93 (SD=24; 46%) trials were relocated. Note that from the valid trials, we discarded those trials that shared a common phase bin in the phase bin acceptance zone, leaving 79 (SD=15) and 86 (SD=13) trials on average for predicted slow and predicted fast trials, respectively. For information on the number of trials in Stage 2 at individual-level, see Table S3.

The mean RT difference across participants between slow and fast phase bins in Stage 2 was −0.439 ms (SD=4), which was not significant according to a group t-test (t(7)=−0.2977, p=0.6127, d_z_=−0.1052). Individually, none of the participants presented a significant difference in RTs between the visual targets presented in the predicted slow and fast phase bins of the α-cycle (all ps > 0.1) (Table 1).

**Table 1.** Individual data for fast/slow phase bins including phase bin acceptance zone in Stage 2. For each participant, the number of trials (max=100), the tested angular points (degrees), and the mean RTs (ms) are reported for the fast and slow phase bins tested in Stage 2. Statistics indicate the results (t value, degrees of freedom, p-value, Cohen’s dz and 95%-confidence intervals CI) of an unpaired t-test (right-tailed, p<0.05) comparing slow vs. fast RTs individually. Group level data and statistics are also reported.

| Part. | Slow phase bin |  |  | Fast phase bin |  |  | RT diff.<br>phases<br>[in ms] | Statistics |  |  |  |  |
| --- | --- | --- | --- | --- | --- | --- | --- | --- | --- | --- | --- | --- |
|  | No.<br>of<br>trials | Phase<br>bin | Mean<br>(SD)<br>RTs<br>[in ms] | No.<br>of<br>trials | Phase<br>bin | Mean<br>(SD)<br>RTs<br>[in ms] |  | t | dof | p | dz | RT diff.<br>95% CI |
| 1 | 88 | 162° | 198 (34) | 85 | 234° | 200 (37) | -2 | -0.30 | 171 | .62 | -.05 | [-10.43<br>7.24] |
| 2 | 100 | 162° | 196 (39) | 100 | 18° | 202 (30) | -6 | -1.26 | 198 | .90 | -.18 | [-14.24<br>1.90] |
| 3 | 74 | 90° | 203 (32) | 59 | 126° | 206 (39) | -3 | -0.54 | 131 | .71 | -.10 | [-11.70<br>5.88] |
| 4 | 66 | 306° | 202 (36) | 86 | 234° | 195 (39) | 8 | 1.24 | 150 | .11 | .20 | [-2.53<br>17.83] |
| 5 | 100 | 54° | 194 (29) | 100 | 306° | 191 (33) | 3 | 0.69 | 198 | .24 | .10 | [-4.18<br>10.27] |
| 6 | 62 | 54° | 207 (34) | 87 | 18° | 207 (32) | 0 | -0.02 | 147 | .51 | -.00 | [-9.05<br>8.85] |
| 7 | 75 | 306° | 200 (32) | 78 | 270° | 201 (40) | -1 | -0.21 | 151 | .58 | -.03 | [-10.97<br>8.50] |
| 8 | 68 | 306° | 191 (39) | 90 | 54° | 193 (37) | -2 | -0.36 | 156 | .64 | -.06 | [-12.07<br>7.68] |
| Mean<br>(SD) | 79<br>(15) | -- | 199 (5) | 86<br>(13) | -- | 199 (6) | 0 (4) | -- | -- | -- | -- | -- |
| Group<br>level | 633 | -- | 199 (5) | 685 | -- | 199 (6) | 0 | -0.30 | 7 | .61 | -.11 | [-5.66<br>4.78] |

### Interim discussion and reality checks

The analyses according to the pre-registered pipeline adapted from Callaway & Yeager (1960) did not return a consistent relation between the phase of individual ongoing α-oscillations and the speed of responses to visual targets at the individual or group level. Compared to offline experimental approaches, where analyses can be adjusted retrospectively, real-time settings imply a priori parameter choices that can affect the outcome. Therefore, we proceeded to exclude the possibility that the null results from the main analyses may have originated from a priori choices in the real-time setting. We focused on three aspects: First, we checked the accuracy of the closed-loop BCI system in sending the stimulus trigger at the intended phases along the α-cycle. Second, we questioned whether the choice of the IFoI based on resting EEG data was representative of the dominant α-frequency during the task. Finally, we checked that the electrode choice (O1) for the real-time α-phase estimation was representative of α-activity of interest in the occipito-parietal cluster of electrodes.

#### Reality check 1: Accuracy of phase estimation during the real-time experiment

The question here was how precisely the closed-loop BCI triggered visual stimuli at the desired phases along the α-cycle. We, therefore, selected all valid trials for each stage and extracted the phase at which the stimulus was presented from the empirical EEG offline, and computed the absolute difference between the desired and the actual phase using the Circular Statistics Toolbox (Berens, 2009). On average, across phase bins and participants, our BCI system hit +1.21° (SD= 33.70°) off the intended phase in Stage 1, and +3.79° (SD= 25.43°) in Stage 2 (see Figure 3 and Figure S2 and S3 for individual results on phase accuracy in Stage 1 and Stage 2, respectively). The phase estimation accuracy of our real-time BCI setting seems comparable to previous attempts at phase-triggered events, like for example targeting the α-frequency in the motor cortex (Zrenner *et al.*, 2018; Hougaard *et al.*, 2019) which typically achieved an accuracy within −12° to 5° off of the desired phase, with SDs between 48° and 55°.

**Figure 3.**
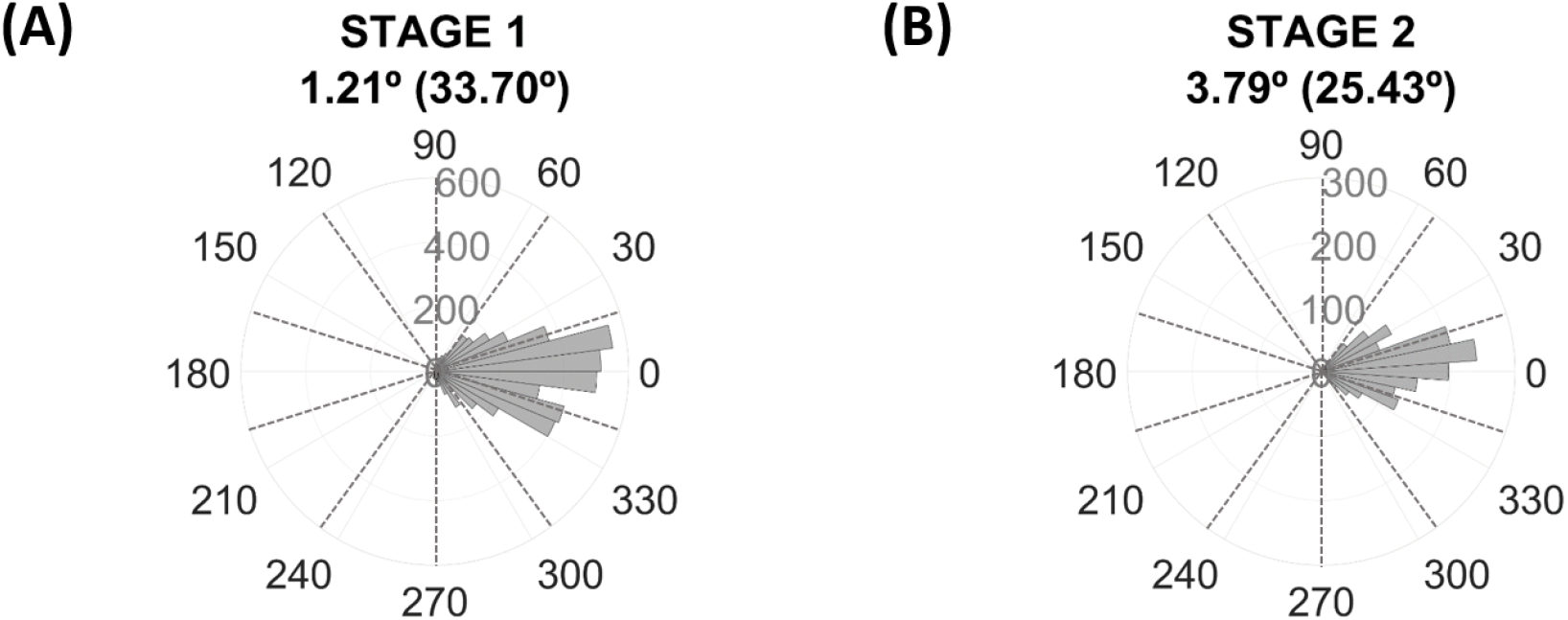
Rose plot of phase accuracy for valid trials across participants **(A)** in Stage 1 (total No. of trials = 4933) and **(B)** in Stage 2 (total No. of trials = 1600). For convenience, all phases have been realigned to 0⁰. Dotted lines denote boundaries between phase bins.

We also decided to check the reasonable expectation of whether the accuracy of the phase estimation depended on the latencies from the reference point of the EEG signal (i.e., phase bins) and whether it was worse at increasing latencies. We rearranged phase accuracy values based on the phase bins and found, as expected, that both the mean and standard deviation of phase accuracy are worse at increasing latencies (see Figure S4 and Table S4 for individual and group data). When subtracting the accuracy of the last-first latencies, the average mean varies −5.42° (SD=9.38°) and the variability increases 32.20° (SD=10.64°).

Finally, we calculated the cumulative percentage of trials as a function of the phase bin difference between target and hit phases. Table S5 shows the results at the individual and group level. Overall, 45% of trials felt in the target phase bin, 88% of trials within ±1 bin, and nearly all trials within ±2 bins.

These overall results show a reliable alignment at each of the targeted phases of the α-cycle. Although extrapolating approximately over one α-cycle has led to more unreliable phase accuracy at increasing latencies, the accuracies are still within safe limits in terms of our purposes. Note that accuracy of phase estimation was checked against real data as part of the BCI setting, so that trials with erroneous estimations that satisfied the trial validity criteria (i.e., RT and amplitude) were eventually relocated if necessary to the hit phase bin in Stage 1 or discarded if did not fall within the acceptance zone in Stage 2. We, therefore, think that the possibility that null results in the α-phase – RTs relationship might be explained by an inaccurate triggering of targets is minimal.

#### Reality check 2: Frequency of interest during the real-time experiment

The real-time stages of the experiment used the IFoI, which was estimated in a 5-min pre-experiment screening session from a cluster of occipito-parietal electrodes (see *Screening and estimation of the IFoI* for more details). This pre-screening is a common practice in order to customize the EEG analysis in terms of individual frequency, especially in the α-band. One potential problem, however, is that the frequency measured in the screening session was not representative of the relevant frequency during the task. Small deviations between the relevant α-frequency during task execution and the one actually used might be mitigated in our protocol because we used a relatively large spectral window (IFoI ± 5 Hz). However, large deviations might affect the subsequent steps of the online protocol, such as filtering and forward prediction. To estimate if such deviations took place, we compared the IFoI used in the real-time experiment stages (that is, the one estimated from the screening stage) against the actual dominant α frequency recorded during task execution. IFoI during task execution was computed following the same procedure as for the IFoI in the screening stage. To avoid potential contamination from visual and motor evoked responses, we used EEG epochs from +500 ms after button press to stimulus onset of the next trial (duration about 2 s, depending on inter-trial jitter). We computed the power spectrum density (PSD) within the α-band (5-15 Hz) for O1-electrode (the electrode in the BCI setting) and for the OP-cluster (the same as used in the screening session for the selection of the IFoI: P7, P3, Pz, P4, P8, O1, Oz, O2) using the Welch method (window=500 ms; overlap=10%; resolution=0.25 Hz). For each participant, the power spectrum was normalized by the mean power in the 1 to 40 Hz window. Figure 4A illustrates the single frequency peak within the α-band during task execution, plotted against the IFoI at rest used in the real-time experiment. Overall, the mean peak difference between rest (OP-cluster) and task (O1-electrode) IFoIs was 0.16 Hz (SD=0.23), with a maximum absolute mean difference of 0.50 Hz in participant 8 (see Table S6 for individual results). IFoI at rest in OP-cluster was very close to the dominant α-frequency during task execution from the same electrode, with deviations of the central frequency of less than 1 Hz.

**Figure 4.**
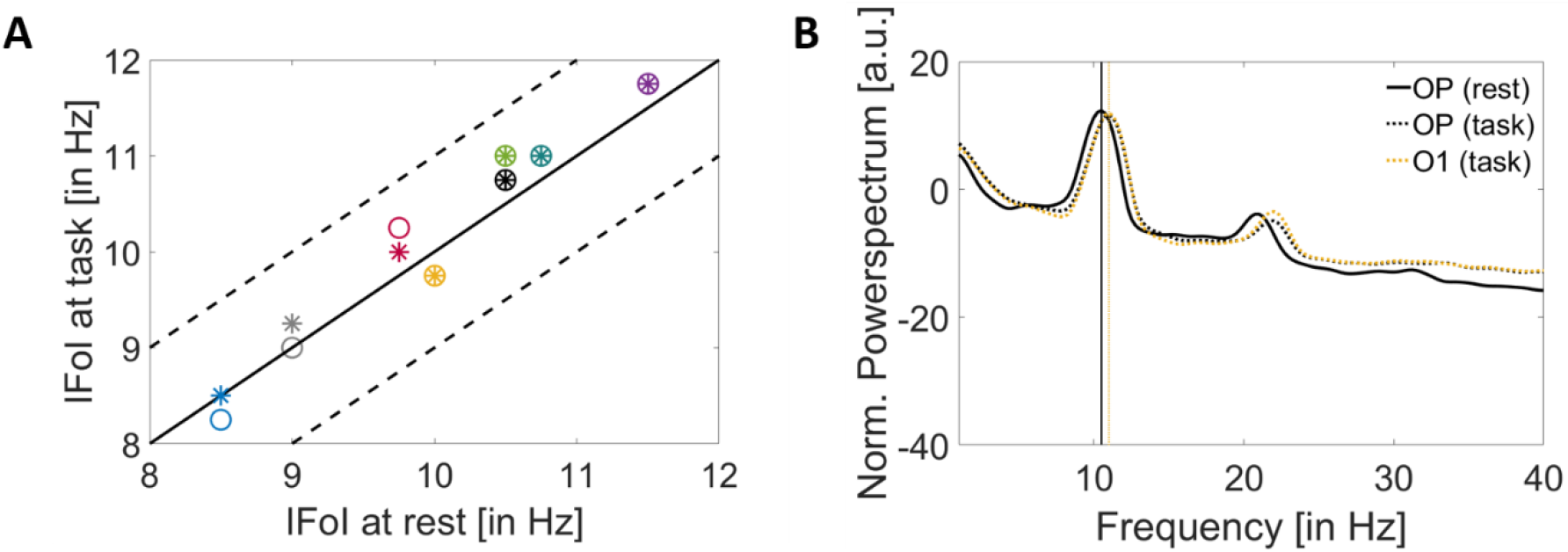
**(A)** IFoI [in Hz] at rest using OP-cluster (P7, P3, Pz, P4, P8, O1, Oz, O2) vs IFoI during the task [in Hz] for two different electrode set conditions: OP-cluster (circles) and O1-electrode (asterisks). Dashed lines denote ±1 Hz and each colour denotes a participant. **(B)** Power spectrum of a representative participant (P08) showing the biggest mean difference (0.50 Hz) between IFoI at rest computed at OP-cluster (solid black line) and IFoI at task computed with O1-electrode (dotted yellow line). Power spectrum computed at OP-cluster at task is also plotted (dotted black line).

Moreover, we decided to check whether the instantaneous frequency differed from the IFoI on a trial-by-trial basis. We calculated the instantaneous frequency for each trial using the Hilbert transform. We band-pass filtered the data from O1 electrode within 5-15 Hz (Butterworth filter order 2, one-pass), epoched from –2 to 2 s (from stimulus onset), demeaned and detrended. We computed the instantaneous frequency using the MATLAB function ‘instfreq’. We selected the prestimulus window of interest of −1 to 0 s from stimulus onset (same time window as the one-second buffer in the real-time experiment) to average the instantaneous frequency within the selected window. Figure S5 shows a heatmap chart of the variation of the phase accuracy as a function of the frequency difference from IFoI for each participant, in which colour denotes the number of trials. All the participants show a mean frequency deviation from IFoI lower than 1 Hz. Participant P08 shows the smaller variability in frequency deviation (mean SD=0.32 Hz), whereas participants P03 and P05 show the larger variability (mean SD=0.73 Hz) (see Table S5). At group-level, the average mean frequency across participants is 0.73 (SD=0.24) Hz.

Note that we were extrapolating approximately one α-cycle from a reference point at the peak of the EEG signal. Therefore, we would expect to see that if the error in the frequency estimation is negative (i.e., actual instantaneous frequency is slower than the estimated IFoI), then there will be a positive error in the phase estimation (i.e., the stimuli would reach an earlier hit phase and the target phase would happen afterwards in time). Hence, we expected the direction of the correlation to be negative. To check for this hypothesis, we decided to compute a linear correlation between the frequency difference from IFoI and the phase accuracy for each participant by adopting a directional one-tailed hypothesis testing. Table S6 shows the results, including Pearson’s coefficient with its p-value. Overall, we see that 5 out of 8 participants present a negative correlation, as expected, and only participants P01 (p=.04), P02 (p=.001) and P07 (p<.001) show a significant negative correlation.

We can, therefore, conclude that the BCI setting employed here was successful in centring the spectral analysis around the desired relevant frequency of interest and variation in frequency from the IFoI at a single-trial level might have influenced the phase accuracy of the BCI setting. However, as stated in the previous section, if there was a deviation in the frequency from the IFoI in a given trial, we took that into account by checking the empirical phase at stimulus presentation and reallocate it if necessary to the hit phase bin in Stage 1 or discarded if did not fall within the acceptance zone in Stage 2.

#### Reality check 3: Electrode of interest used in the real-time experiment

As for the frequency of interest, we had to a priori decide about the electrode of interest to be used in the BCI system. Following Callaway & Yeager (1960), we chose O1-electrode. This choice appeared convenient to limit real-time computational delays due to clustering over a larger set of electrodes. However, it is perhaps important to check that the signal picked up from O1-electrode in the real-time stages was representative of the α-frequency dominant in a wider occipito-parietal cluster. Therefore, we compared the activity in O1-electrode to that of a cluster of occipito-parietal electrodes (OP cluster: P7, P3, Pz, P4, P8, O1, Oz, O2). The spectral comparison was analogous to the one described for the reality check 2 (IFoI rest vs IFoI task). The results, illustrated in Figure 4B (see Figure S6 and Table S6 for individual data), show that the two α estimates were within 0.20 Hz.

In five out of the eight participants, the relevant frequency peak using the occipito-parietal cluster was the same as in O1-electrode. We can, therefore, confirm that the α-fluctuations picked up from O1-electrode as our IFoI during the real-time experiment were closely representative of the occipito-parietal activity.

## EXPLORATORY ANALYSES

In the present study, we aimed at employing a closed-loop BCI approach to show that the phase of ongoing α-oscillations measured with EEG can be harnessed to expedite RTs. This proof-of-concept can not only open avenues for neuro-devices but also help to test the relevance of the α-theories. To do so, we sought to achieve a conceptual replication of a seminal study were such effects had been reported in the past (Callaway & Yeager, 1960). Although our setting included some corrective measures and online checks, we ensured that the system successfully phase-locked visual stimulation to ongoing occipital α-oscillations, the pre-registered analyses returned null results. We decided to explore the data further to find out if phase effects on RTs could be found using other approaches.

### Re-analysis of Stage 2 data including only trials for fast/slow phases

In the main analysis of Stage 2, we decided to include trials in which our online phase estimation was ± 1 phase bins from the intended (fast/slow) phase. This decision was taken under the assumption that, by definition, phase effects fluctuate gradually, so that excitability in phase bins near the maximum peak (minimum peak) would still be relatively high (low). However, we decided to re-do the analysis in Stage 2 and select only those trials falling strictly in the slow and fast phase bins, while excluding those falling in ± 1 phase bin acceptance zone (as well as all the rest, as before). We re-calculated the mean RT between slow and fast phase bins and found a mean difference of −0.625 ms (SD=7), which was not significant neither at individual nor at the group level (t(7)=-0.2496, p=0. 0.5950, dz=-0.0883). Note that the number of trials was much reduced, leaving 55 (SD=18) and 52 (SD=9) trials on average for predicted slow and predicted fast trials, respectively. For information on the number of trials and statistics at individual-level, see Table S7. These findings are in line with those found in the pre-registered results.

### Smoothing the RT-phase modulation

The main purpose of Stage 1 was to estimate the phase bins with fastest and slowest RTs for later use in Stage 2. However, given that we collected a minimum of 50 RTs for each of the 10 bins distributed throughout the α-cycle, one could search for a possible phase-ordered pattern in the RTs in that dataset. As described in the *Results* section, plotting the mean RTs for each phase bin (Figure 2) did not seem to highlight any discernible oscillatory pattern. However, we did not perform a formal statistical analysis at that stage. In this follow up analysis, we adapted an analytical approach used by Fiebelkorn (2013) to test statistically for an oscillatory pattern in Stage 1 data. The logic behind this approach is that if a phase-dependent modulation of RTs exists, then RTs should vary significantly around opposite phases of α (as in the idealized example in Figure 1). Therefore, we looked for pairs of phases 180° apart that could result in a larger difference between RTs. We calculated the average RT over the trials lumped within −90° and +90° around each phase bin to obtain 10 phase-centred RT averages. Then, we normalized each phase-centred RT average by the average RT across all trials. If RTs were modulated by phase, the normalized phase-centred RTs would resemble a sine wave. To test the statistical significance of possible phase modulation, we transformed these phase-centred RT averages to the frequency domain through a fast Fourier Transform (FFT). We tested the significance (α-level = 0.05) of the peak in the FFT for one cycle using a Monte Carlo randomization procedure (10,000 randomizations). The statistical tests showed that none of the participants displayed a significant RT modulation (p-values ranged from 0.11 to 0.92; Figure 5). This outcome confirmed the results from the pre-registered analysis on phase effects, described above.

**Figure 5.**
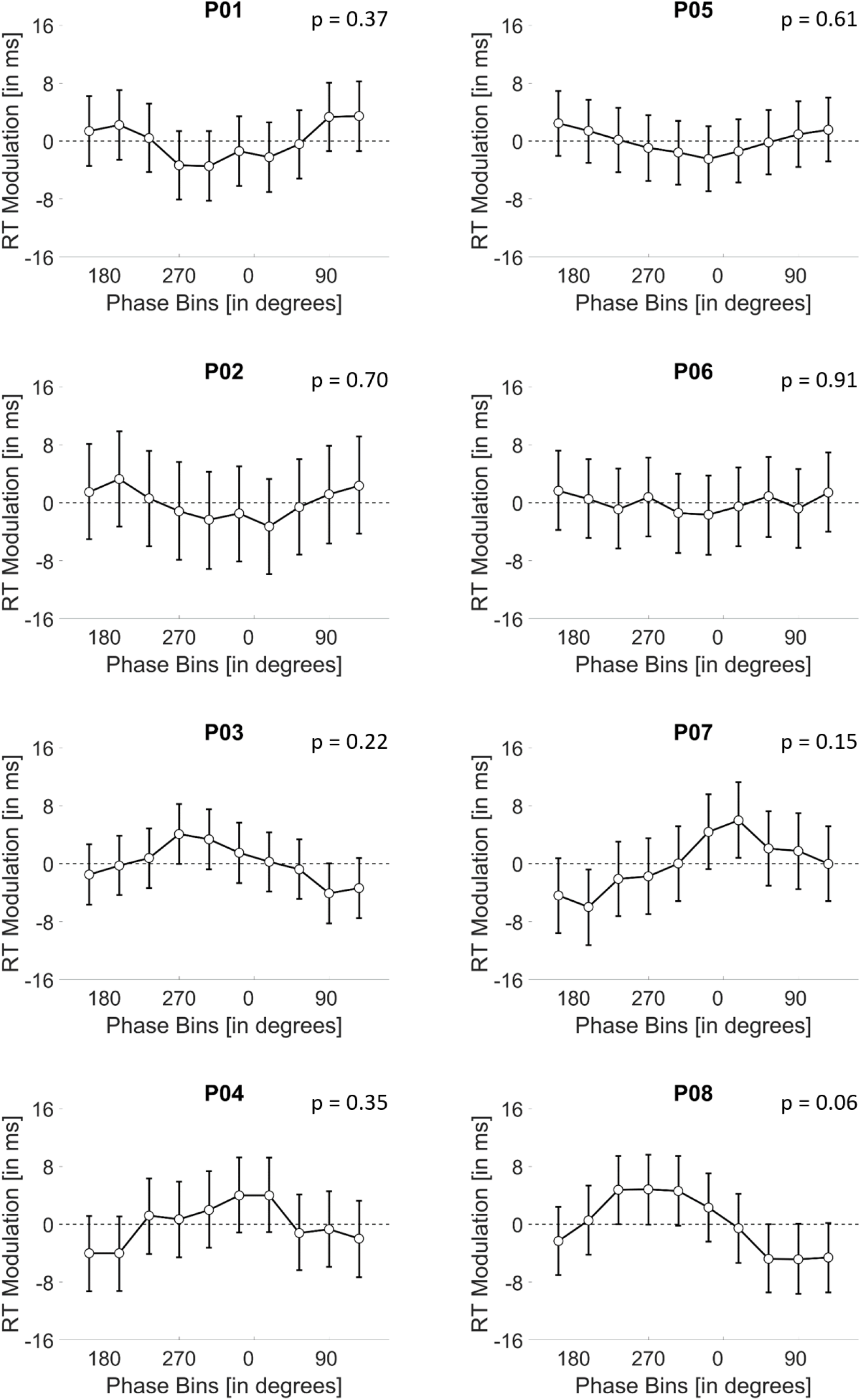
Phase-RT modulation for each phase bin (centred on the target phases) in Stage 1 for all participants. P-value of the modulation comes from using a Monte Carlo randomization procedure (10,000 randomizations). Error bars denote the 95%-confidence intervals (CI) of the randomizations.

### Searching for phase opposition at stimulus onset

If the response to a stimulus is related to the phase of the α-activity, we should expect a pattern of opposition within the α-frequency (the narrow band EEG) when comparing between slow and fast RT trials at stimulus onset (time = 0) (Mathewson *et al.*, 2009). We checked for this possibility by offline band-pass filtering data from electrode O1 using a Butterworth filter (order 2, two-pass) around the IFoI ± 5 Hz band. No re-referencing was applied. All valid trials from Stage 1 were used for the analysis. EEG data were epoched from – 200 ms to 300 ms (t=0 being the stimulus onset time), and then demeaned and baseline corrected (−200 ms to 0 ms). To average across subjects with different IFoI, the time dimension of the EEG was transformed into α-cycle units (the time vector multiplied by the IFoI; data resampled by linear interpolation). RTs were median-split into slow and fast categories. The inter-participants average RT was 231 ms (SD = 9) for slow trials and 182 ms (SD = 8) for fast trials. Figure 6 shows the narrow-band activity for the slow and fast RTs, and for all trials pooled together (see, Figure S7, for individual plots). Although visual inspection suggests an opposition pattern in the narrow-band activity between slow and fast trials at stimulus onset, this pattern is not statistically significant. This can be appreciated by comparing the apparent difference with the large overlap in the confidence intervals (fast trials mean = −0.4454 μV, 95% CI = −1.5599, 0.6691 μV; slow trials mean = 0.3901 μV, 95% CI = −0.6210, 1.4012 μV). However, to test for phase opposition beyond visual inspection, we used the Phase Opposition Sum (POS) method (VanRullen, 2016b). We forward-filtered the data from electrode O1 using a Butterworth filter (order 2, one-pass) around the IFoI ± 5 Hz band and computed the phase by means of the Hilbert transform. We applied the POS on phase values at stimulus onset for fast vs slow trials, both at individual and group-level. The statistical significance was assessed using non-parametric permutations tests (10,000 iterations) using random shuffles of trial assignment to slow/fast bins to compute the distributions of POS values to be expected by chance for each subject. For the group, the distribution of POS values to be expected by chance corresponded to the average of individual POS values to be expected by chance. The p-value associated to measured POS at individual (group) level corresponded to the proportion of times that individual (group averaged) POS obtained in the permutation exceeded measured individual (group averaged) POS. In line with the results of the pre-registered analyses, we did not observe a significant group-level effect of POS between slow and fast RTs (p=.932) nor at individual-level.

**Figure 6.**
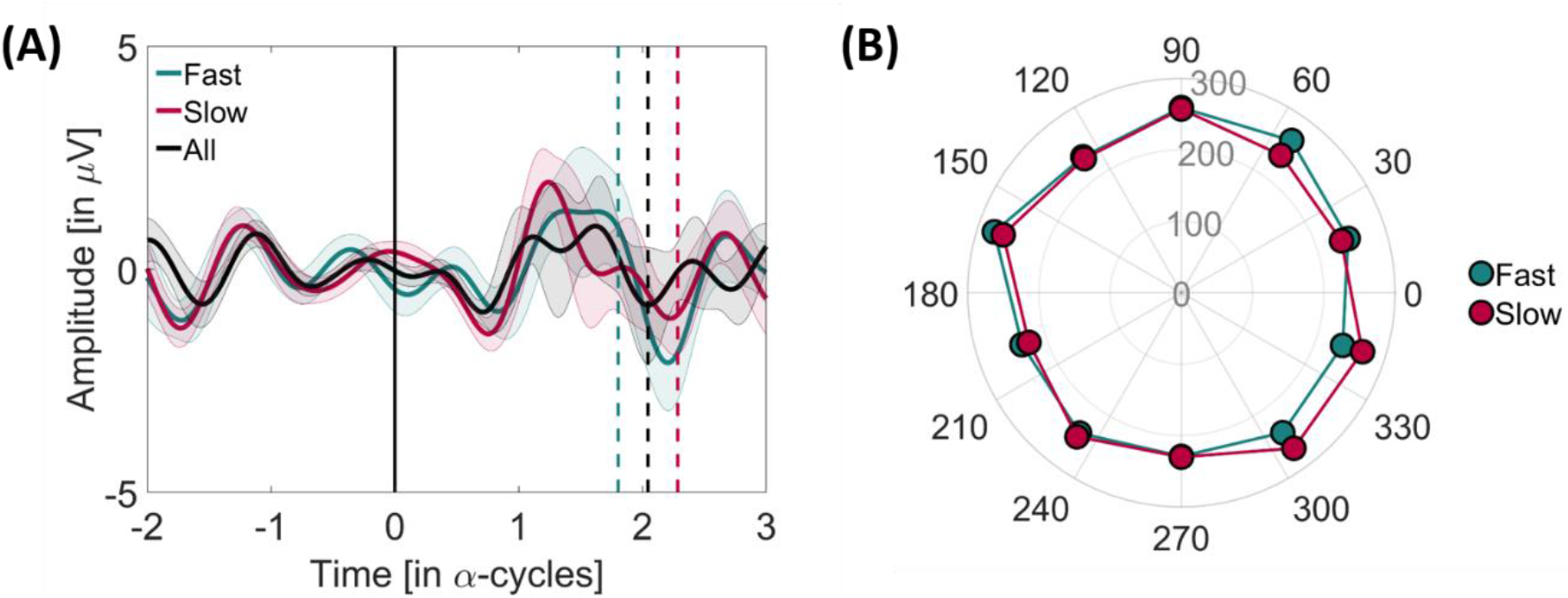
**(A)** Grand average of the narrow-band activity time-locked to visual stimulus presentation for fast (green), slow (red), and all (black) trials from O1-electrode in time of the α-cycles. Thin lines represent the standard error of the mean (SEM) interval. Vertical dashed lines denote the mean RTs for all slow (red), fast (green) and all trials (black). **(B)** Polar plot of the representation of the overall number of trials across participants for each phase bin (dots) for fast and slow trials.

### Effect size equivalence test

The present study was a replication of a previous one testing a relationship between the α-phase and RT (Callaway & Yeager, 1960). The results were at variance with that original study: we did not find evidence for such a relationship. We decided to perform an equivalence testing (Lakens *et al.*, 2018) to compare the effect size in the original study with that reached in the present, even if we had already looked at the data. Callaway & Yeager (1960) reported data from 8 participants and achieved a mean difference between slow and fast RT of 8.13 ms (SD= 5.11; d_z_ = 1.59). In a new study, it would then be reasonable to expect a minimum effect size equal to 33% of the original effect size, assuming the effect in the original study were true (Simonsohn, 2015). This means that we would expect a minimum theoretical effect size of 2.681 ms (or d_z_= 0.53). Incidentally, our aim before the study was to achieve RT differences that would also be relevant in terms of BCI application in real life, and therefore possibly larger than the meagre 2.7 ms effect size derived from the present estimation (admittedly, performed a posteriori). Nevertheless, if not useful at the practical level, one would hope to gather some information at the theoretical level. In the present study, we tested 8 participants and obtained a mean difference between slow and fast RTs of −0.439 ms (SD= 4.171 ms; d_z_ = 0.105) (Stage 2). Given this sample size we cannot reject that the real difference between conditions is bigger than 0 (t(7) = −0.298, p = 0.775, α-level = 0.05, one-tailed) or that the effect size is between −.53 and .53 (t(7) = 1.521, p = 0.09, given equivalence bounds of – 2.68 and 2.68 ms and α-level = 0.05). The equivalence test was ran also at the individual level: None of the participants showed an RT difference in the expected direction over the 2.68 ms limit that it is a reasonable value to be anticipate based on Callaway & Yeager (1960). Note that the objective of this study was to find effects at participant level. Therefore, the group tests performed here and in previous sections are exploratory and must be interpreted with caution as they are very likely underpowered.

## Discussion

The present study aimed at providing a proof of concept for harnessing on the phase of ongoing α-oscillations recorded non-invasively with EEG for real-time BCI, and to garner support for the role of such occipito-parietal α-oscillations in visual perception. Evidence of this kind is valuable because it can help achieve a better understanding of the relation between the occipito-parietal α-phase and behavioural outcome (i.e., speed of reaction times to visual events), and lay the groundwork for possible BCI applications. Our study was a modern replication of Callaway & Yeager’s study (1960), where participants performed a speeded detection on visual targets triggered in real-time at different angles of a participant’s α-cycle. First, we sampled RTs to visual targets presented at 10 different phase bins throughout the α-cycle (Stage 1) to select the phases associated with slowest/fastest RTs. Second, we measured RTs to visual targets presented at these two pre-selected phase bins (Stage 2). If a consistent phase-RT relation exists in the expected direction, it follows that, in Stage 2, stimuli presented at the slow phase would have led to slower RTs compared to stimuli presented at the fast phase.

Contrary to what was expected, the analyses did not return a consistent relation between the phase of ongoing α-oscillations and RTs neither at the group nor at the individual level. Because this experiment was run in real-time, most analytical choices had been made a priori, based on previous literature (Callaway & Yeager, 1960). Please note that obtaining the expected results using a priori set analytical pipeline would implicitly corroborate the brain-behaviour theory behind the decisions for the closed-loop. In this sense, closed-loop BCI can be considered a test bench for brain-behaviour theories. However, because some of the prior choices might have been decisive in producing a null result in the present study, we performed a few reality-checks *a posteriori*. First, we verified that the intended phase of visual stimulation and the actual one were in alignment by comparing the time of stimulus delivery with the empirical EEG measurements, offline. Real-time phase estimation was less than 5° off the intended phase (+1.21±33.70° and +3.79±25.43°, in Stages 1 and 2, respectively), which compares well with estimation accuracy in other modern phase-based closed-loop BCIs (Zrenner *et al.*, 2018; Hougaard *et al.*, 2019). We also demonstrated that the accuracy of the phase estimation depended on the latencies along the α-cycle. We found an average mean variation of −5.42° (SD=9.38°) and an increase of variability of 32.20° (SD=10.64°) between the last and the first latencies. Second, we double-confirmed that the initial choice of the frequency of interest was representative of the predominant α-frequency during task execution by analyzing the data both at single-trial and stage-dataset levels. Third, we validated our choice of the electrode (O1) for the real-time analysis as representative of the central frequency of interest in the occipito-parietal cluster, the most common for the α-rhythm in visual perception (Myers *et al.*, 2014; Samaha *et al.*, 2015; Benwell *et al.*, 2017; Harris *et al.*, 2018; Ruzzoli *et al.*, 2019). Taking all the results together, we reckon that both the estimation over almost one α-cycle and the difference in frequency from IFoI on a trial-by-trials basis are probably the main reasons why phase accuracy varied over latencies along the α-cycle and why we had to reallocate trials in Stage 1 and enlarge Stage 2 acceptance zone to ± 1 phase bin. However, we would like to highlight that, in practical terms, nearly an average of 88% of the trials felt within ±1 bin, which we do not consider to be a poor phase estimation for a BCI setting given the resolution of our EEG system and the method we implemented to estimate the phase by extending a sinus using the IFoI from a reference point in the EEG signal.

It is important to note that even if the data we obtained were variable across participants, the focus of our analysis was on the individual effects because one of the interests in this study was BCI application. The cross-validation protocol implemented in our design (selection and validation of phase bins from Stage 1 to Stage 2 within the same individual) also highlighted a substantial within-participant variability, in many cases leading to opposite trends from Stage 1 to Stage 2 (e.g., the expected fast phase bin returned, on average, the slowest RTs).

As we mentioned in the introduction, the relation between the posterior prestimulus α-phase and behaviour has been (and it still is) based on a popular hypothesis, leading to several sister theories (see Ellingson, 1956, for an overview). For example, the α-phase has been interpreted as a *sensory gateway* (Bartley & Bishop, 1932); a *sensory gateway* with a functional *inhibitory role* (Jensen & Mazaheri, 2010); as a *scanning mechanism* (Walter, 1950); as evidence for *excitability cycles* (Bishop, 1932; Lindsley, 1952; see VanRullen, 2016a, for a similar version). The main point in common between these theories is that reactions (accuracy or RTs) to visual stimuli correlate with (and can be predicted by) the oscillatory activity from the occipito-parietal cortex in the α-band. A critical analysis of the literature shows that this hypothesis has not been free of controversy: Early studies reported no (Walsh, 1952; O’Hare, 1954), or weakly significant effects (Lansing, 1957; Callaway & Yeager, 1960; Callaway, 1962a, 1962b; Dustman & Beck, 1965). Null evidence is also reported in modern times with respect to accuracy (Benwell *et al.*, 2017; Ruzzoli *et al.*, 2019). The present study adds to the previous literature showing that the α-phase/RTs relationship is variable and not reliable when targeted in real-time, at least using extra-cranial EEG.

Perhaps it is worth mentioning at this point that we focused on human non-invasive studies (i.e., EEG or MEG) on the role of the α-phase on perception because one our goals was to provide evidence for the possibility to capitalize on this well-studied relationship for BCI settings. We acknowledge, indeed, that prior studies have frequently found a reliable relationship between α-power and spatial attention (for example Worden *et al.*, 2000; Kelly *et al.*, 2006; Thut *et al.*, 2006) or visual memory (see Palva & Palva, 2007), however, the main focus here was narrowed to visual detection performance (Walsh, 1952; Lansing *et al.*, 1959; Bompas *et al.*, 2015; Benwell *et al.*, 2017; Ruzzoli *et al.*, 2019). In this specific case, despite the α-power/behaviour correlation has been more solidly established in the literature, the present study was not optimized to reveal power/behaviour relationships. Indeed, we introduced measures to achieve a consistently high α-power to facilitate reliable phase estimations for stimulus presentation, resulting in a small variability in α-power^4^.

Returning to the focus of the present study, which relates to the putative effect of α-phase on RTs to visual events, we should consider three critical aspects of our design that might have influenced the negative outcome. First, we asked the participants to perform the task with their eyes closed. The eyes closed strategy, also implemented in Callaway & Yeager’s study (1960), induces higher α-power at occipito-parietal locations which is convenient for reliable phase estimation. However, whether and how performing a perceptual task with the eyes closed jeopardized the outcome is unknown. Excluding the possibility that an eyes-closed condition could also involve sub-cortical generators of the α-activity (Lopes da Silva *et al.*, 1973, but see Bollimunta *et al.*, 2011; Sokoliuk *et al.*, 2019), we did not find any theoretical caveat against the eyes-closed strategy, which was instead technically convenient. Please note that others have successfully used eyes-closed preparations in the past (Lansing *et al.*, 1959; Callaway & Yeager, 1960) and more recently (Lim *et al.*, 2013; Hwang *et al.*, 2015). Based on this, we doubt that the eyes-closed condition may have been critical to producing a null result. The second aspect of our design was to use speeded detection, therefore adopting RTs instead of accuracy as the measure of interest. Even if the α-theories have been related to both (RTs: Walsh, 1952; Lansing, 1957; Lansing *et al.*, 1959; Callaway & Yeager, 1960; Accuracy: van Dijk *et al.*, 2008; Busch *et al.*, 2009), no explicit claims have been made on possible differences between the two measures regarding their sensitivity to prestimulus oscillations. One would believe that if the temporal structure of α-oscillations is important to parse sensory information into perception, then it should be relevant for both RTs and accuracy. Furthermore, unlike accuracy, RT is a continuous measure potentially more sensitive to moment-to-moment variation in excitability than the dichotomic responses in a detection task. Another aspect of our design that merits discussion was that stimulus intensity was supra-threshold and fixed across participants. This is often the approach in experiments measuring RT. Yet, one could perhaps argue that the stimulus was so strong that possibly subtle phase-dependent variations in sensory responses were saturated, thereby having a negligible impact on behaviour. It is difficult to answer this question examining previous literature because luminance levels have been rarely reported in a precise fashion. Callaway (1962a) published the results of a study examining the RT – α-phase relationship with dim and bright visual stimuli (although the actual luminance was not reported) and stated that the depth of such modulation did not vary consistently as a function of brightness. To the best of our knowledge, the only study measuring the RT – α-phase relationship where stimulus intensity was reported clearly is Dustman and Beck (1965). The authors found a consistent effect of RT to α-phase with stimuli of 0.7 lum/m^2^ (that is, 0.128 cd/m^2^) at 40 cm distance to the subject’s (closed) eyes. This is brighter than our stimulus intensity (0.076 cd/m^2^). In the absence of reliable information about the stimulus intensity in past studies, one can also look at the response latencies as a proxy. The average RT in our study was 206 ms (SD=9) in Stage 1, and 199 ms (SD=6) in Stage 2. Past studies where a significant RT – α-phase relationship was reported range from faster responses than ours (167±22 ms, Dustman & Beckman, 1965) to slower (245±15 ms and 236±12 ms respectively for slow and fast RTs in Callaway & Yeager, 1960; 295±51ms and 348±64ms for bright and dim stimuli, respectively in Callaway, 1962a). Therefore, even if one cannot be certain of a possible saturation in neural responses following our stimuli, the stimulus strength and the speed of ensuing latencies were within the range of past studies reporting positive effects.

Finally, a fair question to ask is whether the occipito-parietal α-phase is a critical parameter for perception, but difficult to be extracted from EEG-based closed-loop BCI, or whether it is not critical at all. Fluctuations in neuronal excitability giving way to the oscillatory patterns observable with EEG (and MEG) are ubiquitous in the brain, and the relationship between these fluctuations and neural responses to stimuli is well established in physiology (Bishop, 1932; Buzsáki & Draguhn, 2004). This makes oscillations seen in the EEG appealing candidates to explain and predict behaviour. However, the outcomes of tests regarding the role of occipito-parietal α-phase in the organization of the visual flow of information have been positive (Lansing, 1957; Lansing *et al.*, 1959; Callaway & Yeager, 1960; Mathewson *et al.*, 2009) as well as negative (Walsh, 1952; O’Hare, 1954; Benwell *et al.*, 2017; Ruzzoli *et al.*, 2019). One could argue that a relationship between phase and visual detectability (and hence, response latencies) may exist, but it was obscured by the signal-to-noise variability when recoding from scalp electrodes in EEG. This is, in fact, likely given by the oscillatory patterns in neural excitability so frequently observed in intra-cranial recordings (Bishop, 1932; Lopes Da Silva & Storm Van Leeuwen, 1977; Lakatos *et al.*, 2008) or animal studies (Haegens *et al.*, 2011; Spaak *et al.*, 2012; Fiebelkorn *et al.*, 2018, 2019). Based on this, one would have to conclude that despite the results from the present study are far from significant, they are also not conclusive as to disproof an effect and challenge the α-theories meaningfully.

Apart from theoretical considerations, we also had a second main goal in mind running this experiment: To provide a proof-of-concept for the use of oscillatory phase as a real-time control signal in a BCI. We estimated that a minimum RT difference between slow/fast phases of 2.681 ms could be expected (33% of the effects in Callaway & Yeager, 1960). However, from a more practical perspective, we wonder whether such a small (and variable) effect can be efficiently picked up by scalp EEG and, if so, whether it can be considered meaningful in a BCI application. Saving less than 3 ms in, for example, the efficiency of warning signals would seem close to nothing in most applied contexts.

## Conclusions

Taken together, we must infer that our data do not support a relationship between the phase of α-fluctuations measured extra-cranially and response latencies to visual events. A prudent conclusion is that theoretical and empirical knowledge regarding this relationship may need to progress further to generate enough confidence to attempt the application of the α-theory to neuro-devices. Further research might investigate the influence of parameters such as the eyes-closed strategy, the different sensitivity of discrimination performance vs reaction times as the dependent variable, or the impact of stimulus intensity. We believe that, at present, the effort to implement a closed-loop BCI application based on the relationship of occipito-parietal α-phase measured with EEG and reactions to visual events might not pay off. In addition, we encourage other scientists and BCI practitioners to use BCI settings for hypothesis-testing with a priori set methods in the cognitive neuroscience field as a test-bench for brain-behavioural theories and to explore the feasibility of EEG-based BCI applications.

## Acknowledgements

This research was supported by the *Ministerio de Economia y Competitividad* (PSI2016-75558-P AEI/FEDER to S.S.F.), AGAUR *Generalitat de Catalunya* (2014SGR856 to S.S.F.), Explora Ciencia 2015 (AEI - PSI2015-72568-EXP to M.R.). M.R. was also supported by the *European Commission Individual Fellowship* (Ctrl Code – 794649, H2020-MSCA-IF-2017).

## Conflict of Interest Statement

The authors have no conflict of interest to declare.

## Author Contributions

IVG: Methodology, Software, Validation, Formal analysis, Investigation, Data curation, Writing Original Draft & Editing, Visualization.

LMF: Conceptualization, Methodology, Software, Formal Analysis, Writing – Review & Editing.

MTC: Conceptualization, Writing-Review & Editing, Validation.

MR: Conceptualization, Writing Original Draft & Editing, Supervision, Funding acquisition.

SSF: Conceptualization, Writing Original Draft & Editing, Supervision, Funding acquisition.

## Data Accessibility Statement

Data will be uploaded in Open Science Foundation (OSF) and made public upon publication.

## Abbreviations

BCI: Brain-Computer Interface
RT: Reaction time
MEG: Magnetoencephalography
EEG: Electroencephalography
OSF: Open Science Framework
ITI: Inter-Trial Interval
OP: Occipito-parietal
IFoI: Individual Frequency of Interest
PSD: Power spectrum density
LSL: Lab Streaming Layer
FFT: Fast Fourier Transform
POS: Phase Opposition Sum

## Footnotes

1 We considered that a duration of > 10 min × block (leading to more than 160 minutes approximately of total experimental time + EEG cap montage + debriefing) was unacceptable due to fatigue effects (or sleepiness, easy to happen with eyes closed). These factors can have an impact on the α-activity and, therefore, on our phase estimation in the BCI setting.

2 Please note that it is not possible to infer the proper stimulus intensity from Callaway & Yeager’ (*1960*) experiment; therefore, we chose a fixed arbitrary value that was comfortable for the participants and ensure that response latencies were comparable to those in the original study.

3 We prefer to use the term *Individual Frequency of Interest (IFoI)*, instead of *Individual Alpha Frequency (IAF)*, often used in the literature, because we focused on a frequency range between 5-15 Hz, which spreads out a finer range in the conventional α-band (8-12 Hz).

4 An exploratory analysis, presented in the Supplemental Material, confirms both the low power variability and the null power to RT correlation; see Table S8 and Figures S8 and S9.

## Notes

### Competing Interest Statement

The authors have declared no competing interest.

