## Supplemental Material for "Can the occipital alpha-phase speed up visual detection through a real-time EEG-based brain-computer interface (BCI)?"

### Supplementary Material

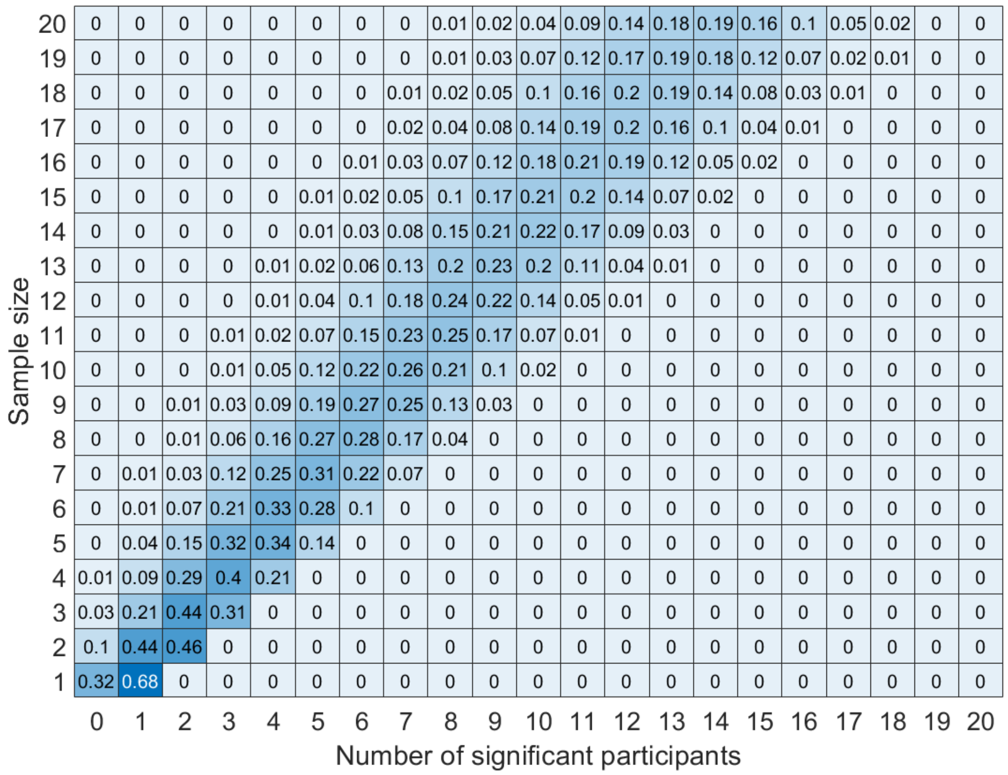


**Figure S1: Sample size estimation**. Based on the results from Callaway and Yeager (1960)'s, we used a Monte Carlo simulation to estimate the probability to find a significant outcome (between RTs at fast and slow phase bins) in a given number of participants (x-axis), depending on the total sample size (y-axis) of our study. The question that our simulation wants to answer was: "*If Callaway's study is representative of the effect in the general population, how probable is to find a significant effect (at participant level) in X participants (x-axis) out of a sample of N participants (y-axis)?*". Each of the cells in the graph estimates the probability of finding X participants with a significant effect when running N participants. According to this simulation, we decided that if less than 3 participants (X<3) out of N=8 showed a significant difference in RTs between fast and slow phase bins, then the size of the effect in this experiment would have to be considered null or negligible compared to the original study (Callaway & Yearger, 1960), assuming an error of 5%.

**Table S1**. Individual results from Stage 1, showing the mean (SD) RT for each phase bin [in ms]. Red and green text represent the slower and the faster RTs for each participant, respectively.

| Part. | Phase bins along α-cycle [in degrees] | | | | | | | | | | Mean (SD) RT combined phases |
|---|---|---|---|---|---|---|---|---|---|---|---|
| | 162° | 198° | 234° | 270° | 306° | 342° | 18° | 54° | 90° | 126° | |
| 1 | **203 (27)** | 196 (32) | **193 (37)** | 196 (23) | 199 (28) | 196 (37) | 196 (35) | 197 (22) | 198 (26) | 199 (24) | **197 (3)** |
| 2 | **203 (38)** | 202 (29) | 194 (46) | 199 (37) | 194 (40) | 203 (40) | **190 (49)** | 196 (45) | 199 (38) | 200 (47) | **198 (4)** |
| 3 | 201 (25) | 207 (29) | 208 (23) | 206 (28) | 201 (28) | 206 (24) | 206 (23) | 202 (26) | **208 (25)** | **200 (23)** | **205 (3)** |
| 4 | 198 (35) | 197 (38) | **193 (27)** | 201 (38) | **210 (35)** | 205 (30) | 198 (34) | 202 (41) | 196 (32) | 196 (30) | **200 (5)** |
| 5 | 199 (26) | 202 (27) | 200 (27) | 201 (25) | **196 (34)** | 204 (25) | 197 (32) | 197 (30) | **204 (30)** | 199 (21) | **200 (3)** |
| 6 | 218 (30) | 221 (32) | 221 (34) | 216 (29) | 217 (33) | 219 (40) | **212 (35)** | 220 (35) | **222 (39)** | 214 (37) | **218 (3)** |
| 7 | 217 (24) | 215 (29) | 215 (38) | **213 (41)** | **230 (30)** | 221 (33) | 217 (32) | 220 (33) | 227 (29) | 218 (27) | **219 (5)** |
| 8 | 207 (25) | 209 (26) | 212 (33) | 207 (36) | **213 (35)** | 208 (35) | 209 (33) | **199 (23)** | 205 (23) | 200 (29) | **207 (5)** |
| **Mean (SD)** | **206 (8)** | **206 (9)** | **205 (11)** | **205 (7)** | **208 (12)** | **208 (8)** | **203 (9)** | **204 (10)** | **207 (11)** | **203 (8)** | **206 (9)** |

**Table S2**. Number of trials in Stage 1. For each participant, it is reported the total number of trials delivered, the number of trials excluded for reaction time (RTs), the number of trials excluded for not satisfying the amplitude threshold criterion, the number of valid trials, the number of hit trials, and the number of trials relocated.

| Part. | No. all trials | No. trials excluded for RTs | No. trials excluded for amplitude | No. valid trials | No. hit trials | No. trials relocated |
|---|---|---|---|---|---|---|
| **1** | **929** | **35** / 927<br>4% | **306** / 927<br>33% | **588** / 927<br>63% | **346** / 588<br>59% | **242** / 588<br>41% |
| **2** | **1259** | **174** / 1259<br>14% | **527** / 1259<br>42% | **558** / 1259<br>44% | **257** / 558<br>46% | **301** / 558<br>54% |
| **3** | **827** | **47** / 827<br>6% | **151** / 827<br>18% | **629** / 827<br>76% | **256** / 629<br>41% | **373** / 629<br>59% |
| **4** | **911** | **76** / 911<br>8% | **183** / 911<br>20% | **652** / 911<br>72% | **244** / 652<br>37% | **408** / 652<br>63% |
| **5** | **1238** | **36** / 1238<br>3% | **542** / 1238<br>44% | **660** / 1238<br>53% | **253** / 660<br>38% | **407** / 660<br>62% |
| **6** | **1194** | **101** / 1194<br>8% | **503** / 1194<br>42% | **590** / 1194<br>50% | **313** / 590<br>53% | **277** / 590<br>47% |
| **7** | **1221** | **122** / 1221<br>10% | **453** / 1221<br>37% | **646** / 1221<br>53% | **277** / 646<br>43% | **369** / 646<br>57% |
| **8** | **695** | **22** / 695<br>3% | **63** / 695<br>9% | **610** / 695<br>88% | **290** / 610<br>48% | **320** / 610<br>52% |
| **Mean (SD)** | **1034 (219)** | **77 (53)**<br>7% | **341 (190)**<br>33% | **617 (36)**<br>60% | **280 (35)**<br>45% | **337 (61)**<br>55% |

**Table S3.** Number of trials in Stage 2. For each participant, it is reported the total number of trials delivered, the number of trials excluded for reaction time (RTs), the number of trials excluded for not satisfying the amplitude threshold criterion, the number of trials excluded for not hitting the phase bin acceptance zone, the number of valid trials, the number of hit trials, and the number of trials relocated.

| Part. | No. all trials | No. trials excluded for RTs | No. trials excluded for amplitude | No. trials excluded for phase accuracy | No. valid trials | No. hit trials | No. trials relocated | No. trials post-hoc excluded |
|---|---|---|---|---|---|---|---|---|
| **1** | **317** | **14** / 317<br>4% | **96** / 317<br>30% | **7** / 317<br>2% | **200** / 317<br>63% | **156** / 200<br>78% | **44** / 200<br>22% | **27** / 200<br>14% |
| **2** | **476** | **58** / 476<br>12% | **196** / 476<br>41% | **22** / 476<br>5% | **200** / 476<br>42% | **116** / 200<br>58% | **84** / 200<br>42% | **0** / 200<br>0% |
| **3** | **440** | **27** / 440<br>6% | **166** / 440<br>38% | **47** / 440<br>11% | **200** / 440<br>45% | **104** / 200<br>52% | **96** / 200<br>48% | **67** / 200<br>34% |
| **4** | **328** | **28** / 328<br>9% | **60** / 328<br>18% | **40** / 328<br>12% | **200** / 328<br>61% | **96** / 200<br>48% | **104** / 200<br>52% | **48** / 200<br>24% |
| **5** | **571** | **33** / 571<br>6% | **267** / 571<br>47% | **71** / 571<br>12% | **200** / 571<br>35% | **73** / 200<br>37% | **127** / 200<br>63% | **0** / 200<br>0% |
| **6** | **494** | **17** / 494<br>3% | **223** / 494<br>45% | **54** / 494<br>11% | **200** / 494<br>40% | **94** / 200<br>47% | **106** / 200<br>53% | **51** / 200<br>26% |
| **7** | **368** | **38** / 368<br>10% | **123** / 368<br>33% | **7** / 368<br>2% | **200** / 368<br>54% | **104** / 200<br>52% | **96** / 200<br>48% | **47** / 200<br>24% |
| **8** | **272** | **35** / 272<br>13% | **22** / 272<br>8% | **15** / 272<br>6% | **200** / 272<br>74% | **115** / 200<br>57% | **85** / 200<br>43% | **42** / 200<br>21% |
| **Mean (SD)** | **408 (103)** | **31 (14)**<br>8% | **144 (84)**<br>35% | **33 (24)**<br>8% | **200 (0)**<br>49% | **107 (24)**<br>54% | **93 (24)**<br>46% | **35 (24)**<br>18% |

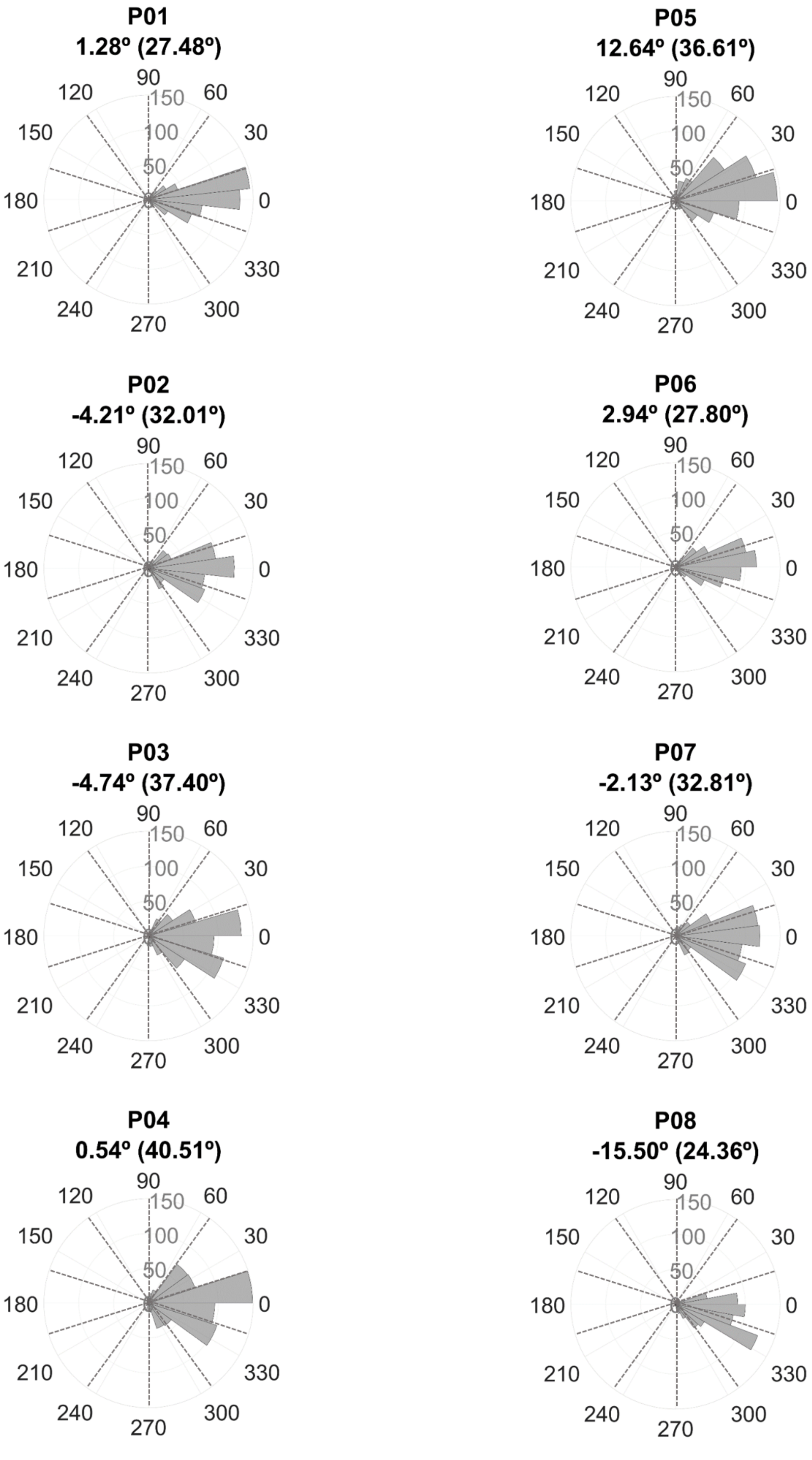


**Figure S2.** Individual Rose plot of mean (SD) phase accuracy of hit phases for all validated trials in Stage 1 [in degrees]. Dotted lines denote boundaries between phase bins.

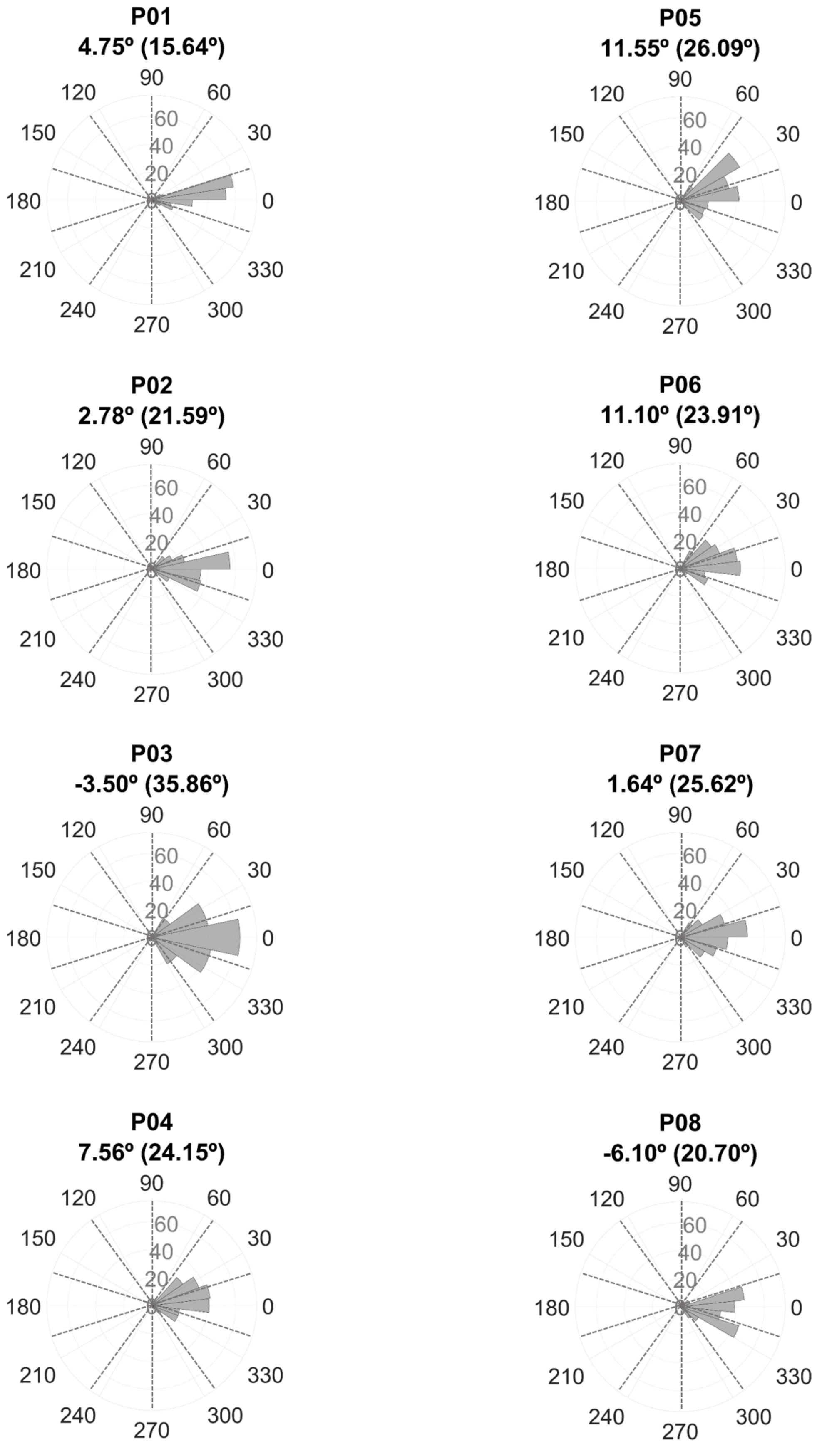


**Figure S3.** Individual Rose plot of mean (SD) phase accuracy of hit phases for all validated trials in Stage 2 [in degrees]. Dotted lines denote boundaries between phase bins.

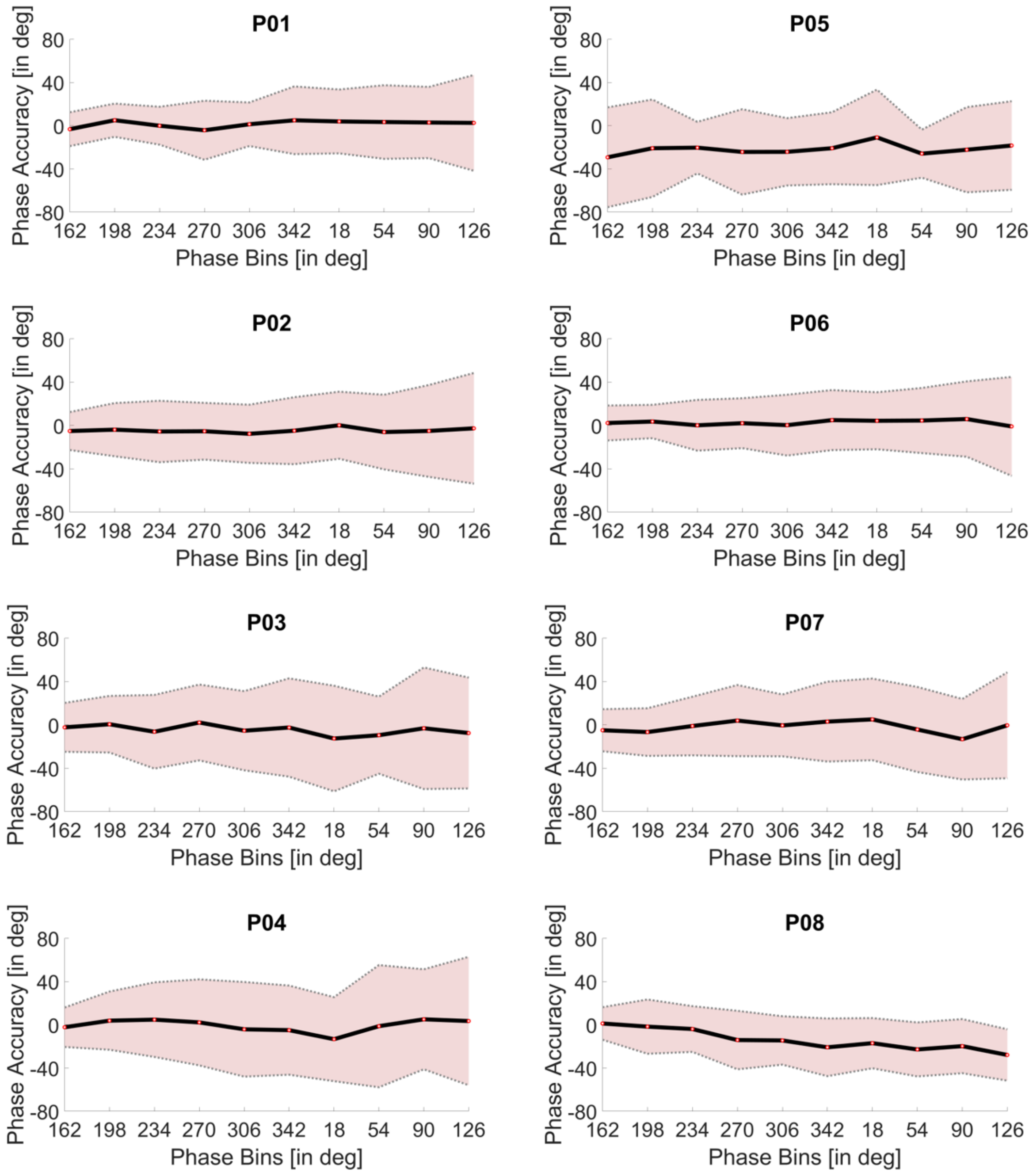


**Figure S4.** Phase accuracy [in degrees] as a function of the phase bins at individual-level. The dark line corresponds to the mean phase accuracy and the shaded area denotes the standard deviation for each phase bin along the α-cycle.

**Table S4.** Difference in mean and standard deviation accuracy computed between the last (i.e., phase bin 126º) and first (i.e., phase bin 162º) phase bins along the α-cycle at individual and group level in Stage 1.

| **Part.** | **Δmean last-first bins [in deg]** | **Δstd last-first bins [in deg]** |
|---|---|---|
| **1** | 5.81 | 28.64 |
| **2** | 2.51 | 33.51 |
| **3** | -5.25 | 28.57 |
| **4** | 5.74 | 41.15 |
| **5** | 10.91 | -5.20 |
| **6** | -3.16 | 29.52 |
| **7** | 4.54 | 29.43 |
| **8** | -29.10 | 8.71 |
| **Mean (SD)** | **-5.42 (9.38)** | **32.20 (10.64)** |

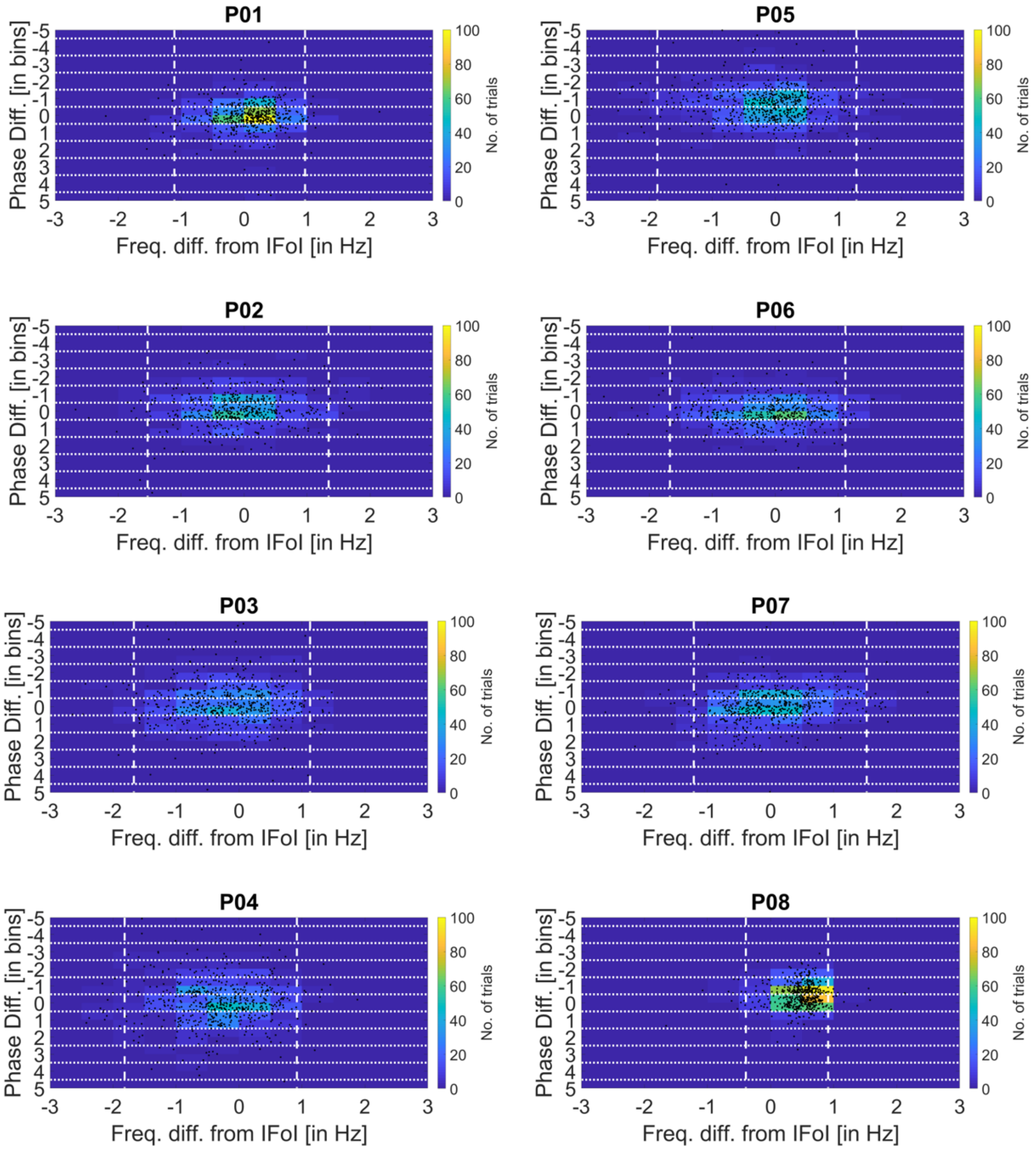


**Figure S5.** Heatmap chart of phase difference [in bins] as a function of the frequency difference of valid trials compared to IFoI at individual-level. Vertical dashed lines denote the interval comprising 95% of the trials, and horizontal dotted lines correspond to phase bin boundaries. The colorbar counts the number of trials and black dots correspond to trials.

**Table S5.** Comparison of the cumulative percentage of trials for phase bin difference between target and hit phase bins. Mean and standard deviation of the frequency difference between Individual Frequency of Interest (IFoI) peak [in Hz] and mean instantaneous frequency [in Hz] during task for Stage 1. Correlation (negative direction, one-tailed) between phase accuracy and frequency difference from IFoI peak and mean instantaneous frequency for Stage 1, including Pearson's coefficient (r) and p-value (p).

| **Part.** | **Cumulative percentage of trials for phase bin difference (target/hit)** | | | | | | **Mean (SD) freq. diff. from IFoI [in Hz]** | **Correlation between phase acc. and freq. diff. from IFoI** | |
|---|---|---|---|---|---|---|---|---|---|
| | **0** | **±1** | **±2** | **±3** | **±4** | **±5** | | **r** | **p** |
| 1 | 59% | 93% | 98% | 99% | 100% | 100% | 0.07 (0.50) | -0.07 | .043 |
| 2 | 47% | 88% | 97% | 99% | 100% | 100% | -0.11 (0.70) | -0.13 | .001 |
| 3 | 40% | 84% | 95% | 98% | 99% | 100% | -0.19 (0.73) | -0.01 | .37 |
| 4 | 37% | 81% | 94% | 98% | 100% | 100% | -0.31 (0.70) | 0.00 | .52 |
| 5 | 35% | 83% | 96% | 98% | 100% | 100% | -0.13 (0.73) | 0.09 | .99 |
| 6 | 53% | 93% | 99% | 100% | 100% | 100% | -0.14 (0.71) | 0.03 | .79 |
| 7 | 44% | 87% | 98% | 99% | 100% | 100% | 0.03 (0.70) | -0.24 | <.001 |
| 8 | 47% | 92% | 100% | 100% | 100% | 100% | 0.48 (0.32) | -0.04 | .15 |
| **Mean (SD)** | **45 (8)** | **88 (5)** | **97 (2)** | **99 (1)** | **100 (0)** | **100 (0)** | **-0.04 (0.24)** | **--** | **--** |

**Table S6.** Comparison among the Individual Frequency of Interest (IFoI) peak [in Hz] and amplitude [in dB] at rest and during task execution for each participant using the occipital-parietal cluster (OP-cluster) and O1-electrode. The IFoI difference is computed between rest (OP-cluster) and task (O1-electrode).

| Part. | IFoI (rest) | | IFoI (task) | | | | ΔIFoI rest-task [in Hz] |
|---|---|---|---|---|---|---|---|
| | OP-cluster | | OP-cluster | | O1-electrode | | |
| | Peak [in Hz] | Amplitude [in dB] | Peak [in Hz] | Amplitude [in dB] | Peak [in Hz] | Amplitude [in dB] | |
| 1 | 10.50 | 10.50 | 10.75 | 5.19 | 10.75 | 10.93 | -0.25 |
| 2 | 11.75 | 6.54 | 11.75 | -8.37 | 11.75 | 7.25 | 0 |
| 3 | 9.75 | 8.00 | 10.25 | 5.28 | 10.00 | 8.24 | -0.25 |
| 4 | 9.00 | 9.99 | 9.00 | 1.19 | 9.25 | 9.12 | -0.25 |
| 5 | 9.75 | 8.06 | 9.75 | 0.80 | 9.75 | 3.02 | 0 |
| 6 | 10.75 | 5.23 | 11.00 | 6.76 | 11.00 | 8.38 | -0.25 |
| 7 | 8.50 | 9.06 | 8.25 | -2.16 | 8.50 | 8.78 | 0 |
| 8 | 10.50 | 12.33 | 11.00 | 11.68 | 11.00 | 11.92 | -0.50 |
| **Mean (SD)** | **10.06 (1.03)** | **8.71 (2.26)** | **10.22 (1.16)** | **2.55 (6.12)** | **10.25 (1.07)** | **8.46 (2.67)** | **-0.19 (0.18)** |

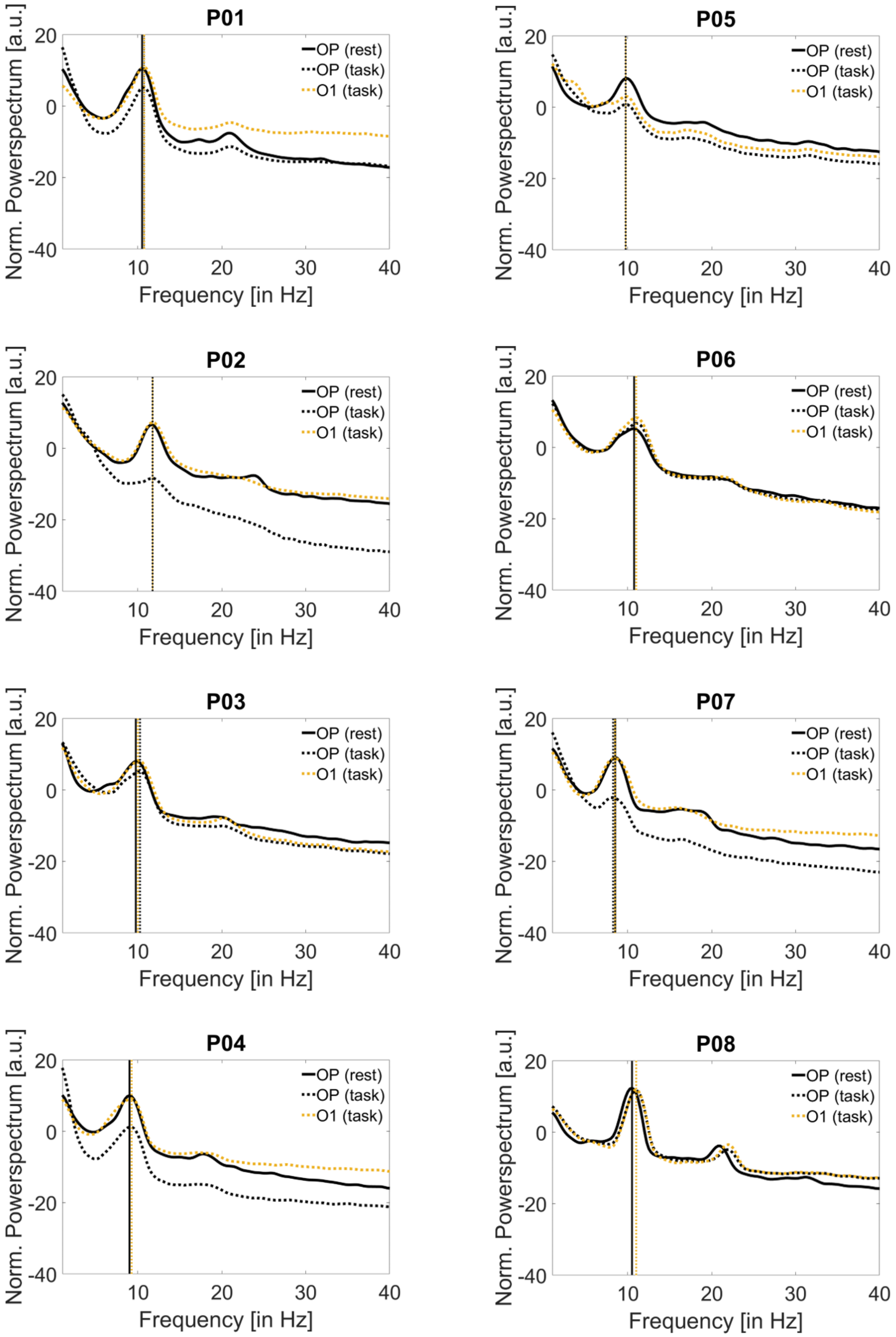


**Figure S6**. Normalized power spectrum for each participant at rest (5-min eyes-closed recording) and during the task (Stage 1 dataset) from 1 to 40 Hz computed with OP-cluster and O1-electrode.

**Table S7.** Individual data for fast/slow phase bins in Stage 2 including only trials that hit the fast/slow phases. For each participant, the number of trials (max=100), the tested angular points (degrees), and the mean RTs (ms) are reported for the fast and slow phase bins tested in Stage 2. Statistics indicate the results (t value, degrees of freedom, p-value, Cohen's $d_z$ and 95%-confidence intervals CI) of an unpaired t-test (right-tailed, $p<0.05$) comparing slow vs. fast RTs individually. Group level data and statistics are also reported.

| | Slow phase bin | | | Fast phase bin | | | | Statistics | | | | |
|---|---|---|---|---|---|---|---|---|---|---|---|---|
| **Part.** | **No. of trials** | **Phase bin** | **Mean (SD) RTs [in ms]** | **No. of trials** | **Phase bin** | **Mean (SD) RTs [in ms]** | **RT diff. phases [in ms]** | **t** | **dof** | **p** | **dz** | **RT diff. 95% CI** |
| 1 | 85 | 162° | 198 (34) | 71 | 234° | 201 (37) | -3 | -0.51 | 154 | .69 | -.08 | [-12.22 6.44] |
| 2 | 68 | 162° | 201 (39) | 48 | 18° | 210 (31) | -9 | -1.25 | 114 | .89 | -.24 | [-19.77 2.75] |
| 3 | 60 | 90° | 199 (31) | 44 | 126° | 203 (27) | -3 | -0.53 | 102 | .70 | -.11 | [-12.76 6.54] |
| 4 | 47 | 306° | 204 (37) | 49 | 234° | 194 (42) | 10 | 1.28 | 94 | .10 | .26 | [-3.04 23.68] |
| 5 | 29 | 54° | 189 (35) | 44 | 306° | 201 (27) | -3 | -0.46 | 71 | .68 | -.11 | [-15.24 8.53] |
| 6 | 40 | 54° | 209 (36) | 54 | 18° | 201 (27) | 8 | 1.30 | 92 | .10 | .26 | [-2.29 19.07] |
| 7 | 48 | 306° | 202 (30) | 56 | 270° | 201 (43) | 1 | 0.07 | 102 | .47 | .01 | [-11.60 12.66] |
| 8 | 64 | 306° | 190 (39) | 51 | 54° | 196 (32) | -6 | -0.90 | 113 | .81 | -.17 | [-17.28 5.07] |
| **Mean (SD)** | **55 (18)** | **--** | **199 (7)** | **52 (9)** | **--** | **200 (5)** | **-1 (7)** | **--** | **--** | **--** | **--** | **--** |
| **Group level** | **441** | **--** | **199 (7)** | **417** | **--** | **200 (5)** | **-1** | **-0.25** | **7** | **.60** | **-.09** | **[-6.38 5.20]** |

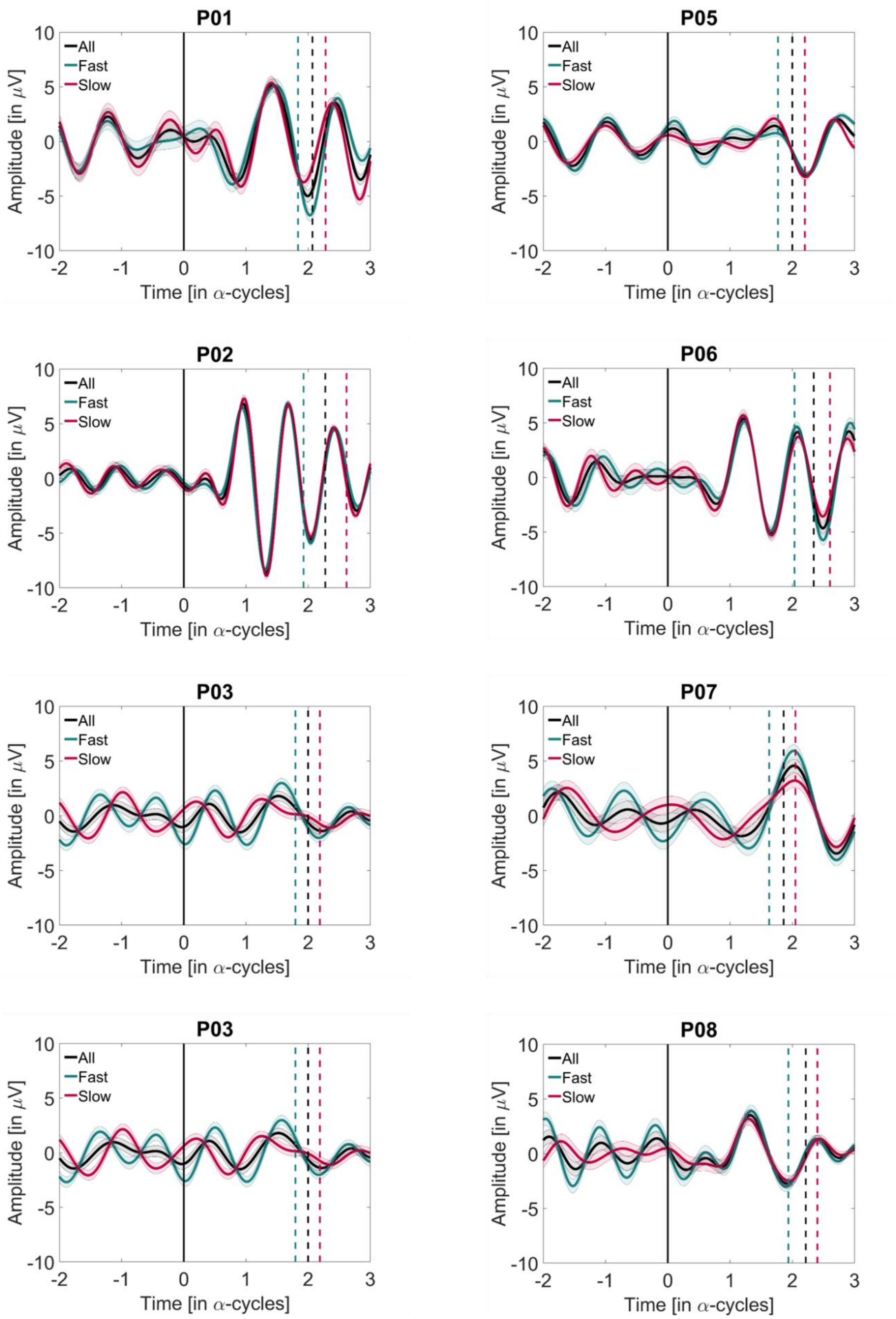


**Figure S7.** Individual narrow-band ERPs for subjects from Stage 1 dataset time-locked to visual stimulus presentation denoting valid trials (black), fast trials (green), and slow trials (red). All plots include SEM interval shading. Dashed vertical lines are plotted to show the mean RT of the trials belonging to each subject (Supporting Table 1).

*Correlation between RTs and prestimulus α-power*

According to α-theories, in addition to phase dependency, the power of spontaneous pre-stimulus oscillations in the α-band has relevance for perception (Lindsley, 1952; Lansing *et al.*, 1959; Klimesch *et al.*, 2007b; Mathewson *et al.*, 2011; Jensen *et al.*, 2014). Despite the fact that our approach was not optimized to find behavioural differences as a function of power, we decided to explore a possible effect of pre-stimulus α-power in relation to slow/fast RTs, as suggested by previous evidence (Walsh, 1952; Lansing *et al.*, 1959; Bompas *et al.*, 2015; Ruzzoli *et al.*, 2019). Specifically, we anticipated that low pre-stimulus α-power would be associated with faster RTs. To this aim, we selected trials from Stage 1 (excluding the RT criterion 50 - 300 ms, in order to increase RT variability). To maximize potential differences in RTs as a function of the α-power, for each participant, we calculated the pre-stimulus power associated with the lower and the higher terciles of the RTs distribution. Data were band-pass filtered 5-15 Hz (Butterworth filter order 2, two-pass), epoched from –1000 ms to 0 ms (i.e., stimulus onset), demeaned and detrended. We computed power by means of the Hilbert transform and averaged within the epoch and across the occipito-parietal cluster used to calculate the IFoI. Please note that this cluster of electrodes has been previously associated to power-behavioral modulation (Myers *et al.*, 2014; Bompas *et al.*, 2015; Samaha *et al.*, 2015; Benwell *et al.*, 2017; Harris *et al.*, 2018; Ruzzoli *et al.*, 2019). Power was normalized with respect to the mean power of each participant and transformed to decibels. Group-level data (see Figure S6 and Table S6 for individual results, and Figure S7 for group results) were analysed using a t-test (left-tailed, $\alpha = 0.05$). No relationship was detected between pre-stimulus α-power and RTs at group ($t(7)= -0.0059$, $p = 0.4977$, $d_z = -0.0021$) nor at the individual level (all ps > 0.1) for all but one single participant ($t(584)= -4.1441$, $p = 0.00002$, $d_z= -0.3424$).

We speculate that the null relation between α-power and RTs in these data is caused by a lack of variability. Please note that despite this relationship has been reliably established in the literature (Walsh, 1952; Lansing *et al.*, 1959; Bompas *et al.*, 2015; Benwell *et al.*, 2017; Ruzzoli *et al.*, 2019), the present study was not optimized to reveal it. In particular, there was little variability in α-power during the experiment, because we introduced measures to achieve a consistently high α-power throughout the task (e.g., eyes closed) to facilitate reliable phase estimations. To ascertain this possibility, we compared the variability in α-power in the present data with the data of another experiment where an effect of power has been found on a visual unspeeded detection task (Ruzzoli *et al.*, 2019, data can be found here: https://osf.io/adrwv/). We found that α-power variability was about 4.5 times lower in the present study compared to the previous one (SD = 2dB vs. SD = 9 dB, respectively).

**Table S8**. Number of trials, mean (SD) RT, and mean (SD) log-transformed and normalized pre-stimulus α-power (dB) for slow and fast trials. Individual t-tests assess statistical difference in the α-power ($p<0.05$). Only P01 showed a significance difference in the α-power between slow and fast trials.

| | Slow trials | | | Fast trials | | | Statistics | |
|---|---|---|---|---|---|---|---|---|
| **Part.** | **No. of trials** | **Mean (SD) RTs [in ms]** | **Mean (SD) ΔPower [in dB]** | **No. of trials** | **Mean (SD) RTs [in ms]** | **Mean (SD) ΔPower [in dB]** | **t** | **p =** |
| **1*** | 293 | 226 (20) | 0.04 (0.94) | 293 | 167 (20) | -0.35 (1.30) | -4.14 | .00002 |
| **2** | 336 | 238 (24) | -0.16 (1.25) | 336 | 156 (28) | -0.16 (1.27) | -0.02 | .49 |
| **3** | 257 | 234 (20) | -0.15 (1.02) | 257 | 179 (11) | -0.08 (1.06) | 0.83 | .80 |
| **4** | 273 | 239 (25) | -0.13 (0.92) | 273 | 167 (16) | -0.06 (0.95) | 0.87 | .81 |
| **5** | 392 | 228 (21) | -0.40 (1.47) | 392 | 170 (17) | -0.25 (1.45) | 1.38 | .92 |
| **6** | 341 | 256 (19) | -0.21 (1.14) | 341 | 182 (16) | -0.06 (1.08) | 1.75 | .96 |
| **7** | 346 | 251 (19) | -0.24 (1.29) | 346 | 181 (28) | -0.14 (1.23) | 0.97 | .83 |
| **8** | 225 | 238 (20) | -0.06 (1.23) | 225 | 175 (21) | -0.20 (1.16) | -1.28 | .10 |
| **Mean (SD)** | ***308 (55)*** | ***239 (22)*** | **-0.12 (1.16)** | ***308 (55)*** | **172 (20)** | **-0.16 (1.19)** | **--** | **--** |

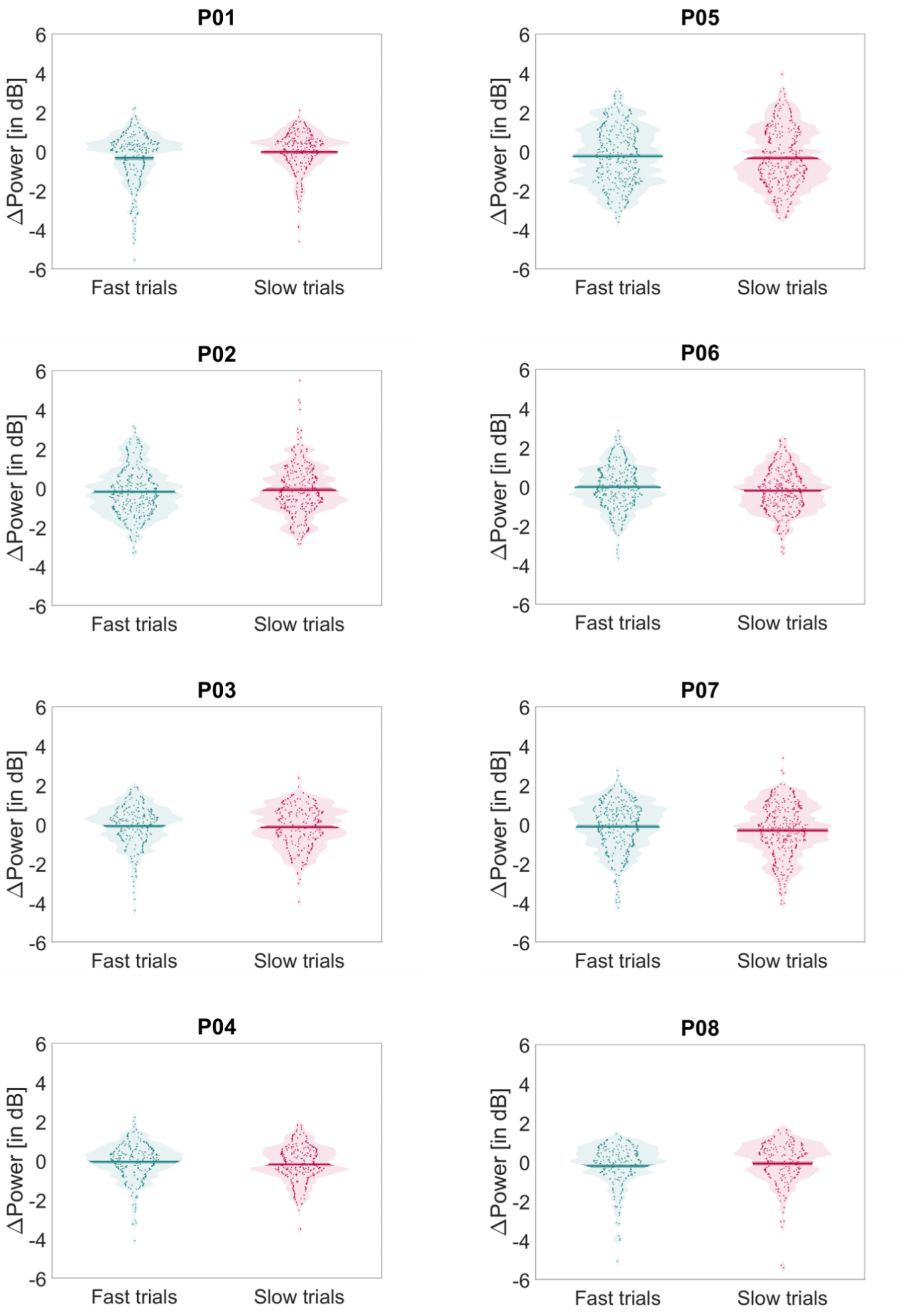


**Figure S8**. Individual pre-stimulus α-power log-transformed and normalized by the mean power [in dB] over all trials in Stage 1 for fast (green) and slow (red) trials. Dots denote the power for each trial at individual level. Violin plots include the mean ± SEM power at individual level.

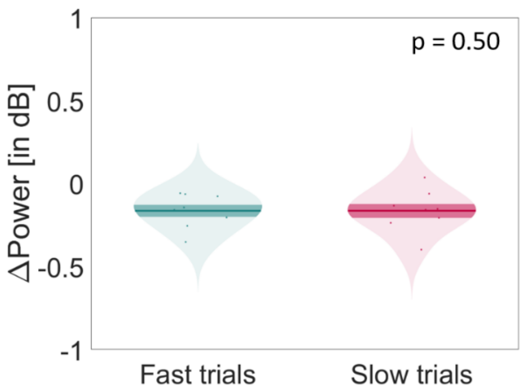


**Figure S9.** Mean (dark line) ± SEM (shaded area) pre-stimulus α-power at group-level; dots denote normalized mean power at individual-level, y-axis represents normalized power [in dB] for the fast and slow trials (x-axis). P-value of the result of the paired-test of the mean power from fast/slow trials.
